# CryoGrid-PIXUL-RNA: High throughput RNA isolation platform for tissue transcript analysis

**DOI:** 10.1101/2022.04.01.486750

**Authors:** Scott A. Schactler, Stephen J. Scheuerman, Andrea Lius, William A. Altemeier, Dowon An, Thomas J. Matula, Michal Mikula, Maria Kulecka, Oleg Denisenko, Daniel Mar, Karol Bomsztyk

## Abstract

Disease molecular complexity requires high throughput workflows to map disease pathways through analysis of vast tissue repositories. Great progress has been made in life sciences analytical technologies. To match the high throughput of these advanced analytical platforms, we have previously developed a multipurpose microplate sonicator, PIXUL, that can be used in multiple workflows to extract analytes from cultured cells and tissue fragments for various downstream molecular assays. And yet, the sample preparation devices, such as PIXUL, along with the downstream analytical capabilities have not been fully exploited to interrogate tissues because storing and sampling of such specimens remain, in comparison, inefficient. To mitigate this bottleneck, we have developed a low-cost user-friendly system, the CryoGrid, that consists of CryoBlock, thermometer/thermocouple, and QR coded CryoTrays to freeze and store frozen tissue fragments, and hand-held CryoCore tool for tissue sampling supported by iPad and Google apps to display tissues, direct coring and share metadata.

RNA is one of the most studied analytes. There is a decades-long history of developing methods to isolate and analyze RNA. Still, the throughput of sampling and RNA extraction from tissues has not matched that of the high throughput transcriptome analytical platforms. To address this need, we have integrated the CryoGrid system with PIXUL-based methods to isolate RNA for gene-specific qPCR and genome-wide transcript analyses. TRIzol is commonly used to isolate RNA but it is labor-intensive, hazardous, requires fume-hoods, and is an expensive reagent. We developed a PIXUL-based TRIzol-free RNA isolation fast protocol that uses a buffer containing proteinase K (PK). Virtually every disease (and often therapeutic agents’ toxicity) is a systemic syndrome but often only one organ is examined. CryoGrid-PIXUL, integrated with either TRIzol or PK buffer RNA isolation protocols, yielded similar RNA profiles in a multiorgan (brain, heart, kidney and liver) mouse model of sepsis. Thus, RNA isolation using the CryoGrid-PIXUL system combined with the PK buffer offers an inexpensive user-friendly workflow to study transcriptional responses in tissues in health and disease as well as in therapeutic interventions.

## INTRODUCTION

We have previously developed a microplate sonicator, PIXUL, that offers unparalleled sample preparation throughput capabilities for a broad range of high throughput analytical applications (1–3). In sharp contrast, available tissue storing and sampling tools lack the throughput to fully exploit the PIXUL sample preparation capabilities to interrogate solid biospecimens.

Freezing is a common way to preserve tissues for storage and transport (4–6). Typically, tissues are snap-frozen using either liquid nitrogen or dry ice/isopentane and stored in vials. Space shortages in deep freezers (−80°C) are a recurring problem for many labs, especially those that process hundreds of tissue samples. Unfortunately, and not infrequently, older samples are discarded to make room for new samples. Laboratory freezers may contain thousands of samples in tubes marked with handwritten or printed numbers, dates, and sample types. Tubes are stored in small cardboard boxes that are also labeled and are either stored loose or are kept in sliding drawers in freezer racks. Without a map guided by codes, finding specific samples can be a challenge and typically involves pulling the racks out one at a time until the needed box is found. Biopsy needles have been used for decades to examine tissue pathology. These are designed for soft tissues, and their through length is 10-20mm. As such, traditional biopsy devices are not suitable to sample frozen tissues that are hard. There are also punch needles that could be used to sample frozen tissue, but it is difficult to get consistent core sizes and to remove frozen cores from the needle’s tip. For a host of molecular analysis and histology, it would be advantageous for researchers, as well as clinical labs, to freeze/store solid tissue specimens in such a way that the same piece of frozen tissue is suitable for multiple samplings, without thawing. To match the high throughput sample preparation power of PIXUL and downstream analytical methods, we developed the CryoGrid system for cryostoring and sampling tissues.

Friedrich Miescher’s isolation and discovery of DNA in 1871 was revolutionary and ushered in the field of nucleic acids research (7). Interestingly, on the 150th anniversary of his discovery, it was shown that the historic nucleic acids isolated by Miescher, in addition to DNA, also contained RNA (8). Then it took decades of work before chemical and biological differences between DNA and RNA began to emerge (9). More than fifty years ago, Temin (10) and Baltimore (11) discovered reverse transcriptase, which allowed the synthesis of complementary DNA (cDNA) from an RNA template. Thirty-five years ago, Mullis discovered polymerase chain reaction, PCR (12,13). On this background of breakthrough discoveries, in 1987 Chomczynski and Sacchi published a single-step method for isolation of RNA (14,15), accelerating the field of RNA biology. Since then, many improvements and iterations of RNA isolation from a variety of substrates have been made including single cells (16–19). There are high throughput RNA, DNA, and protein extraction commercially available instruments and kits. And yet, they require the use of homogenized tissues prior to analyte extraction that slows down the process. Here we developed and tested the CryoGrid system integrated with the PIXUL sonicator for high throughput RNA isolation from tissues for RNA-qPCR and RNA-seq analyses.

## MATERIALS

Hardware/labware (**Table S1**) and kits/enzymes (**Table S2**) catalog numbers and commercial suppliers are listed in the supplementary tables.

### Reagents

TRIzol (Life Technologies 15596018). O.C.T. compound (Fisher Scientific 4585). Dithiothreitol (DTT, D0632), EDTA (E3134), Tris–HCl (T3253) were from Sigma. Sodium chloride (NaCl S-271-3) and Triton X-100 (BP151) from Fisher Scientific. NP40 (198596) from MP Biomedicals. Dulbecco’s Modified Eagle Medium (DMEM-SH30021.0) from HyClone, penicillin/streptomycin (P/S 15749) from Invitrogen, fetal bovine serum (FBS 43635-500) from Jr. Scientific, and phosphate buffered saline (PBS 70013-032), Chloroform (J.T. Baker, 9180-01), Ethanol (Decon Labs, 2716), and Isopropanol (Acros Organics,3223-0010).

### Buffers

Preparations of all buffers and stock solutions were done with nuclease free reagents and ultrapure distilled RNAse/DNAse free H_2_O. PBS: 137 mM NaCl, 10 mM Sodium phosphate, 2.7 mM KCl, pH 7.4; TE: 10 mM Tris-HCl, 1 mM EDTA, pH 7.5; Immunoprecipitation (IP) buffer: 150 mM NaCl, 50 mM Tris–HCl (pH 7.5), 5 mM EDTA, NP-40 (0.5% vol/vol), Triton X-100 (1.0% vol/vol); PIXUL-RNA PK buffer: 10mM Tris-HCl pH 8.0, 10mM EDTA, 0.5% SDS, 50μg/ml Proteinase K, and 40mM DTT.

## METHODS

### Mouse sepsis model and organ harvesting

Female 8–12-week-old C57bl/6 mice were injected intraperitoneally (IP) with 5mg/kg lipopolysaccharide (LPS) in 200μl PBS or with 200μl PBS as control. The specific protocol was approved by the Institutional Animal Care and Use Committee (IACUC) at the University of Washington. After 12 hours mice were euthanized by isoflurane overdose and confirmatory cervical dislocation. Brains, hearts, kidneys, and livers were harvested, immediately placed on ice, and then frozen in CryoTrays (see below). Organs from four LPS and four PBS (control) treated mice were used in RNA RT-qPCR and RNA-seq.

### Cell Lines and Treatment

Human kidney HEK293 cell lines were grown in DMEM supplemented with glutamine, penicillin, streptomycin, and 10% fetal bovine serum in round-bottom 96-well polystyrene plates at density ~200,000 cells per well. For time-point experiments, cells were serum-deprived (0.1% FBS) overnight and at specific time points were treated with either 10% FBS as we described previously (1).

### Matrix quantitative reverse transcription real time PCR (Matrix RT-qPCR)

Isolated RNA was reverse transcribed with Superscript, 0.2 mM dNTP (GeneScript, 95040-880), and random hexamers in 10μl reactions in 96-well microplates for 10 minutes at 50°C then 10 minutes at 80°C. RT reactions were diluted 10-fold with elution buffer prior to running qPCR. Housekeeping genes were used to normalize qPCR results (20). RT-qPCR primers are listed in supplementary **Table S3**. We used our previously developed software, PCRCrunch, to acquire, store and analyze qPCR data sets generated by Matrix RT-qPCR (21).

### mRNA-sequencing (RNA-seq)

After isolation, RNA was run through Zymo RNA Clean & Concentrator. Sequencing libraries were prepared using Zymo-Seq RiboFree Total RNA Library Kit with RNA between 205-480ng and libraries amplified between 13-14 cycles of PCR as per manufacturer’s protocol. Quality of libraries was assessed by Agilent 4200 TapeStation system, qPCR with organ and sepsis specific primers (to show retained specificity) and Collibri Library Quantification Kit. Libraries were diluted as per Illumina protocol to a final pooled loading concentration of 650pM in resuspension buffer (RSB) plus Tween 20 with a 10% PhiX spike-in and sequenced in Illumina P2 cartridges on NextSeq 2000 that employed a dual-index, paired-end, 61 base read length (PE61).

Quality control was done for all sequencing fastq.gz files with FastQC (22). Visualization of read coverage was done using the RSeQC ‘geneBody_coverage.py’ function with default settings(23). FastQC and RSeQC figures and data were compiled for visualization with MultiQC (24).

Reads were mapped to the NCBI Genome Reference Consortium Mouse Build 39 (GRCm39) RefSeq assembly. Reads were aligned and counted in RStudio using the Bioconductor ‘RSubread’ (version 2.8.2) package (25). Alignment was done with the ‘align’ function on the gzFASTQ files creating sorted BAM files. ‘featureCounts’ function was used with the NCBI GRCm39 RefSeq Annotation and the commands: isGTFAnnotationFile=TRUE, countMultiMappingReads = FALSE, strandSpecific = 2, isPairedEnd=TRUE.

Differential gene expression (DGE) was done in RStudio using the Bioconductor ‘EdgeR’(26–28) (version 3.36.0) package using a quasi-likelihood F test and GLM approach to do pairwise comparisons between groups. For the QL model, buffer type (TRIzol or PK buffer), organ type, and treatment (LPS vs PBS) were used as factors. Differentially expressed genes were considered significant if the false discovery rate (FDR) was below 0.05. DGE figures were made with Bioconductor ‘EdgeR’, ‘Gli,’a (29)’ and ‘Vidger’ packages (30).

Principal component analysis was performed in RStudio with the Bioconductor ‘EdgeR’ package. The ‘plotMDS.DGEList’ function was used on the filtered and normalized EdgeR DGElist data with method=“LogFC”, gene.selection = “common”, and using the top 500 genes.

The overrepresentation of DEGs within the Reactome (31) pathways was performed with R package lusterProfiler (version 3.6), separately for up- and downregulated DEGs (32). A heatmap visualization was conducted with the ComplexHeatmap R package (version 2.3.1) (33), using pathways identified by both TRIzol and PK buffer conditions and with a ratio of at least 0.1 of differential vs. nondifferential genes present in a pathway. For clarity, only the top 2 pathways from each Reactome Level 1 category for each data point were taken into account. Pearson’s correlation coefficient for DEGs was calculated with R package Hmisc (34).

Use of RSeQC requires .bam files created using the ‘samtools’ software kit so a separate set of bam files was created using a pipeline of: ‘TrimGalore’ (https://www.bioinformatics.babraham.ac.uk/projects/trim_galore/, version 0.6.7) (to trim adapters on the paired reads, ‘hisat2’ (version 2.2.1) with “--rna-strandness RF” to align sequences to the mm10 assembly(35), and samtools (version 1.13) for generation of sorted and indexed bam files(36).

## RESULTS AND DISCUSSION

### CryoGrid system

The complexity and heterogeneity of disease pathways require analysis of large numbers of tissue samples. Almost any disease, and often therapeutic intervention, are systemic conditions where animal models provide the means to understanding multiorgan dysfunction. To store and sample large numbers of biospecimens for multi-omic analysis, we designed a platform for freezing and cryostoring multiple tissue samples and engineered a hand-held rotary tool for rapid sampling of frozen tissues (**Fig.1 and S1–S2**). This system, which we call the CryoGrid, consists of CryoBox, CryoBlock, thermometer/thermocouple, QR barcoded CryoTrays and CryoCore. The CryoBox is a Styrofoam box filled with dry ice pellets. There is a small hole in the wall of the box to pass a thermocouple wire. CryoBlock, machined from aluminum, with 24 (4×6) cube-sized pockets in the top surface to accommodate 4×6 CryoTray (**Fig.S1**), is seated in the dry-ice-containing CryoBox. The top surface of the CryoBlock is tilted 30° to optimize ergonomics. The CryoBlock has a small aperture on the side, close to the block top, to insert the tip of the thermocouple probe to monitor the block’s freezing and maintenance temperature. CryoBlock chilled in the CryoBox (<−70°C) is used to freeze tissues in the CryoTrays and/or keep the samples frozen while extracting cores.

**Fig.1.**
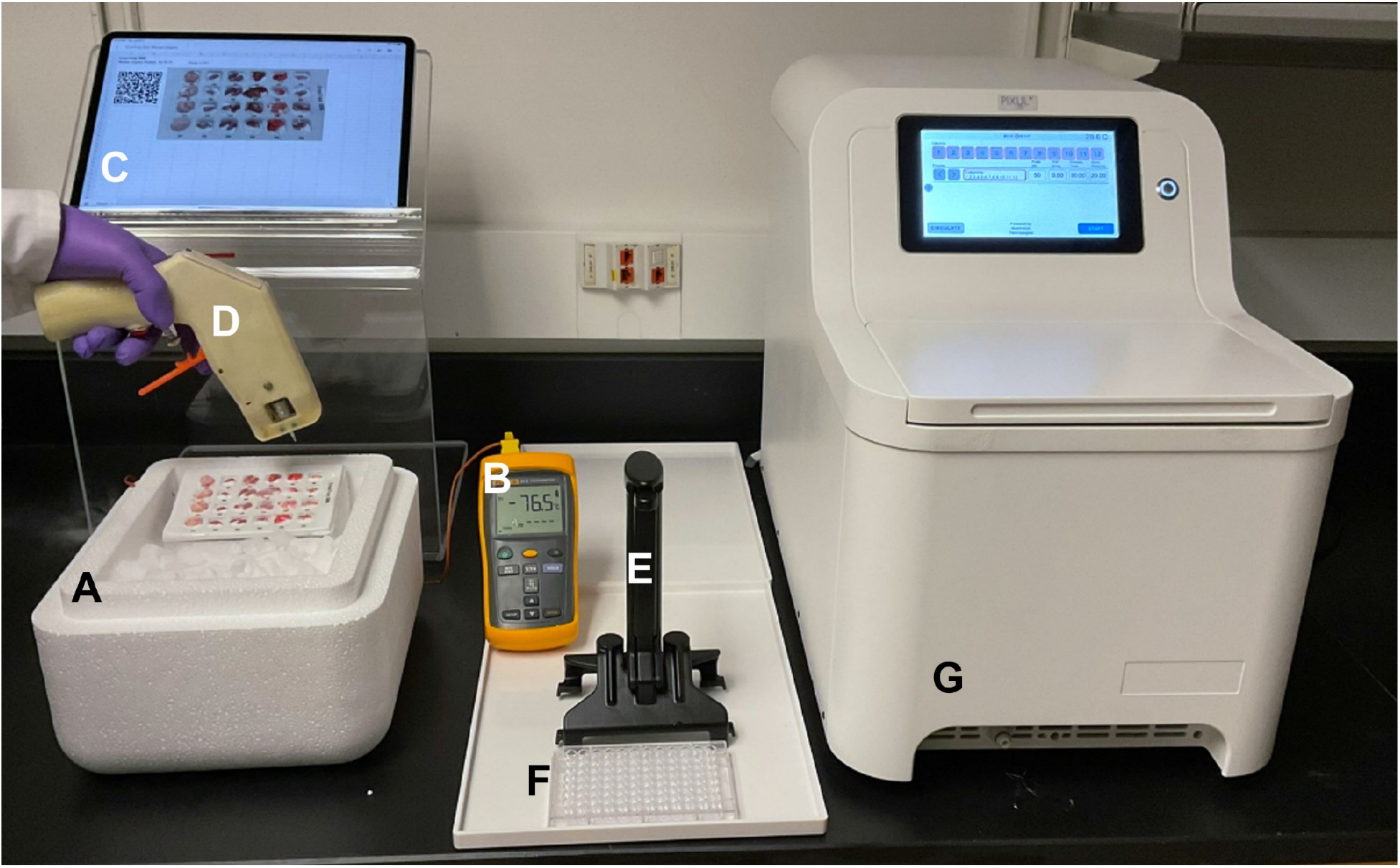
CryoGrid-PIXUL system. ***A***, CryoTray (with QR code recorded online in Google Drive, see Fig.1S) is seated in a CryoBlock cooled in an off-the-shelf Styrofoam box containing dry ice pellets (CryoBox)(< −70°C). Tissue pieces are placed in CryoTray pockets following a pre-designed experimental template in Google Drive. Tissues are frozen by adding optimal cutting temperature (OCT) or CryoGel cryogenic media. CryoTray-embedded frozen tissue layout is then photographed (iPad) and uploaded online to the matching QR-coded Google Sheet containing relevant metadata. ***B***, Thermometer thermocouple probe wire is threaded through a hole at the side of a CryoBox and inserted into an aperture on the side of the CryoBlock to monitor the temperature. ***C***, iPad, on an acrylic stand, displays the CryoTray layout of annotated tissues, provides the means to read or hear recorded notes, and to type-in and/or dictate comments using voice recording in online Google Sheet. ***D***, CryoCore is a motor-driven miniature hole-saw tool to extract cores of frozen tissues embedded in CryoGrid and then eject the cores directly into wells of 96-well PIXUL plates. ***E***, PlateHandle is a tool to facilitate the transfer of 96-well plates in-and-out of PIXUL. ***F***, 96-well plate for sonicating samples in PIXUL. ***G***, PIXUL instrument.

CryoTrays are heat-molded from polystyrene sheets into rectangular trays that contain an array (6×4 wells) of round corner cube-shaped (1×1×1cm) pockets that serve as receptacles for freezing and storing tissues (**Figs.S1–S2**). CryoTrays are covered with a transparent polystyrene lid that has a QR code that uniquely identifies a given CryoTray. The QR code is entered into a web-based Google Drive storage database allowing users to enter, edit and share metadata of the tissues frozen in each QR-coded CryoTray. The CryoTrays with frozen tissues identified with a QR-code are stored in a deep freezer in standard pull-out aluminum drawers at −80°C. An upright deep freezer can store >100,000 ~1gm frozen tissue fragments in 24-wells CryoTrays.

To engineer a novel-design hand-held battery-powered rotary tool for rapid sampling of frozen tissues, CryoCore (**Fig.1 and S1**), we used the idea of a trephine (from Greek word trypanon, an instrument for boring). With CryoCore, the same frozen tissue can be sampled multiple times without thawing, yielding reproducible core sizes (~1-2mm^3^ or ~1-2 mg). The CryoCore design was tailored for multisampling tissues stored in CryoTrays (**Fig.S2**) The estimated (by DNA content) number of cells/tissue core are as follows: brain ~0.4 e+06; heart ~0.3 e+06; kidney ~1.2 e+06 and liver ~0.9 e+06.

### Tissue freezing, sampling, jetting cores into PIXUL plate and sonicating

Before freezing tissues, a 24-well CryoTray is placed into the chilled CryoBlock maintained at < −70°C in a CryoBox filled with dry ice pellets. After harvesting, mouse organs are immediately put on ice and then one by one are placed in individual pockets of the 24 wells and small amounts of embedding matrix (e.g., OCT or Leica CryoGel) are injected into the wells for rapid freezing and immobilizing of tissue fragments (**Fig.1**). The total amount of time to freeze 24 mouse organs is less than 20 minutes. The CryoTrays with the organs covered by a QR code labeled lid is stored at −80°C.

For sampling, the CryoTray with frozen organs is inserted in a chilled CryoBlock (< −70°C) in a CryoBox filled with dry ice pellets (**Fig.1**). An iPad is used to display a Google Sheet document with the organ layout legend and convenient access to metadata to facilitate sampling with the CryoCore. Before coring, the CryoCore trephine is cooled by plunging it in the dry ice pellets. CryoCore tissue cores are jetted with PBS (drawn from the CryoCore syringe reservoir) directly into wells of a 96-well round-bottom heat-resistant polypropylene PIXUL plate kept on ice. The 96-well plate is covered with a MicroAmp Optical Adhesive tape and a small “V” is cut in the top of each well to allow jetting samples into the wells while preventing cross-contamination. One or two cores /well are sampled from each tissue fragment. After collecting all the samples, the plate is covered with a replacement optical adhesive tape, centrifuged for 30 sec at 500xg, the optical adhesive tape is then removed carefully and discarded, and the PBS buffer from CryoCore sample ejection is aspirated from the wells. 100μl of either PK buffer or TRIzol are added to the wells, and the 96-well plate is covered with a new optical adhesive film and placed into the PIXUL sonicator (Matchstick Technologies, Inc Kirkland WA and Active Motif Carlsbad, CA). Samples are processed for 30 seconds in PIXUL with settings: Pulse=50, PRF=1.0 Burst=20. The 96-well plate with sonicated samples is centrifuged for 30 seconds at 500 ×g to collect debris at the bottom of the wells.

### RNA isolation using PIXUL

TRIzol is a commonly used reagent to isolate RNA (16) but the protocol is labor-intensive, relatively slow and uses hazardous solvents requiring working in safety hoods. Further, TRIzol reagent is expensive and involves costly material shipment. We set out to develop a faster, lower-cost protocol without the use of hazardous materials that can be done on the bench. Proteinase K (named for its ability to hydrolyze keratin (37)) is a broad-spectrum serine protease which proteolytically inactivates nucleases even in the presence of sodium dodecyl sulfate (SDS) (37–39). There is a long history of using this enzyme to isolate nucleic acids from biospecimens without the need to use hazardous solvents (38–41). We have previously formulated proteinase K containing buffer to elute DNA from immunoprecipated chromatin (42,43). In cell culture experiments we used serum-deprived human kidney HEK293 cells that were activated with serum treatment leading to a transient induction of immediate early genes such as *EGR1*(*1*). PIXUL multisample sonicator allows one to isolate analytes from 96-well plate cultures of HEK293 cells without sample transfer greatly facilitating protocol develoment (1).

Proteins with high binding activity for ribonucleases (RNase) potently inhibit RNases (44) such as the commercially available recombinant RNaseOUT (**Table S2**). Serum-deprived HEK293 cells were treated with 10% serum for 0, 30, 60, and 120 minutes, supernatants were aspirated, and either proteinase K-elution buffer, with or with RNaseOUT, or TRIzol was added to individual wells, and plates were sonicated in PIXUL. The proteinase K samples were transferred to tubes, boiled, put on ice, centrifuged and the supernatants were collected. TRIzol samples were used to purify RNA using a standard under the safety hood protocol. Contaminating genomic DNA (gDNA) in RNA extraction is a challenge, especially for analysis of rare transcripts requiring the use of DNases to remove even minor amounts of gDNA (45,46). To minimize contaminating gDNA, the RNA prepared using each protocol was treated with or without DNase I (**Table S2**) and then used in reverse transcriptase (RT) reaction to generate cDNAs as templates in real-time PCR (qPCR) with primers to serum-inducible and housekeeping genes. The results showed the predicted serum-induced transient increase in *EGR1* expression (1) and the increase was greater in DNase I treated samples with either the proteinase K or TRIzol method (**Fig. S3**). **Fig. 2** shows the qPCR data as cycle threshold, CT (lower CT values correspond to higher cDNA amount (47), CT lower by 1.0 is equal to 2-fold higher DNA levels), using primers that span (*EGR1*: Exon1-Exon2; *NR1A1:* Exon7-Exon8) or not (*EGR1*: Exon2; *NR1A1*: Exon7) exon regions. There was a marked transient serum-induced decrease in CTs for the *EGR1* and *NR4A1* primers spanning exons in the DNase I treated samples. In contrast, with the *NR4A1* Exon7 primers without DNase I treatment both proteinase K and TRIzol samples had lower CTs (22 vs 30-32) and failed to show serum induction indicating gDNA contamination. The use of PCR primers that span exons does not completely overcome the misreading of the mRNA levels, particularly for the less abundant transcripts such as *NR4A1*, and underscores the fact that even with TRIzol DNase I treatment appears to be essential for preparing RNA used in RT-qPCR analysis. In this protocol, the results suggest that the use of RNaseOUT did not make a difference in extracting RNA from HEK293 cells.

**Fig.2.**
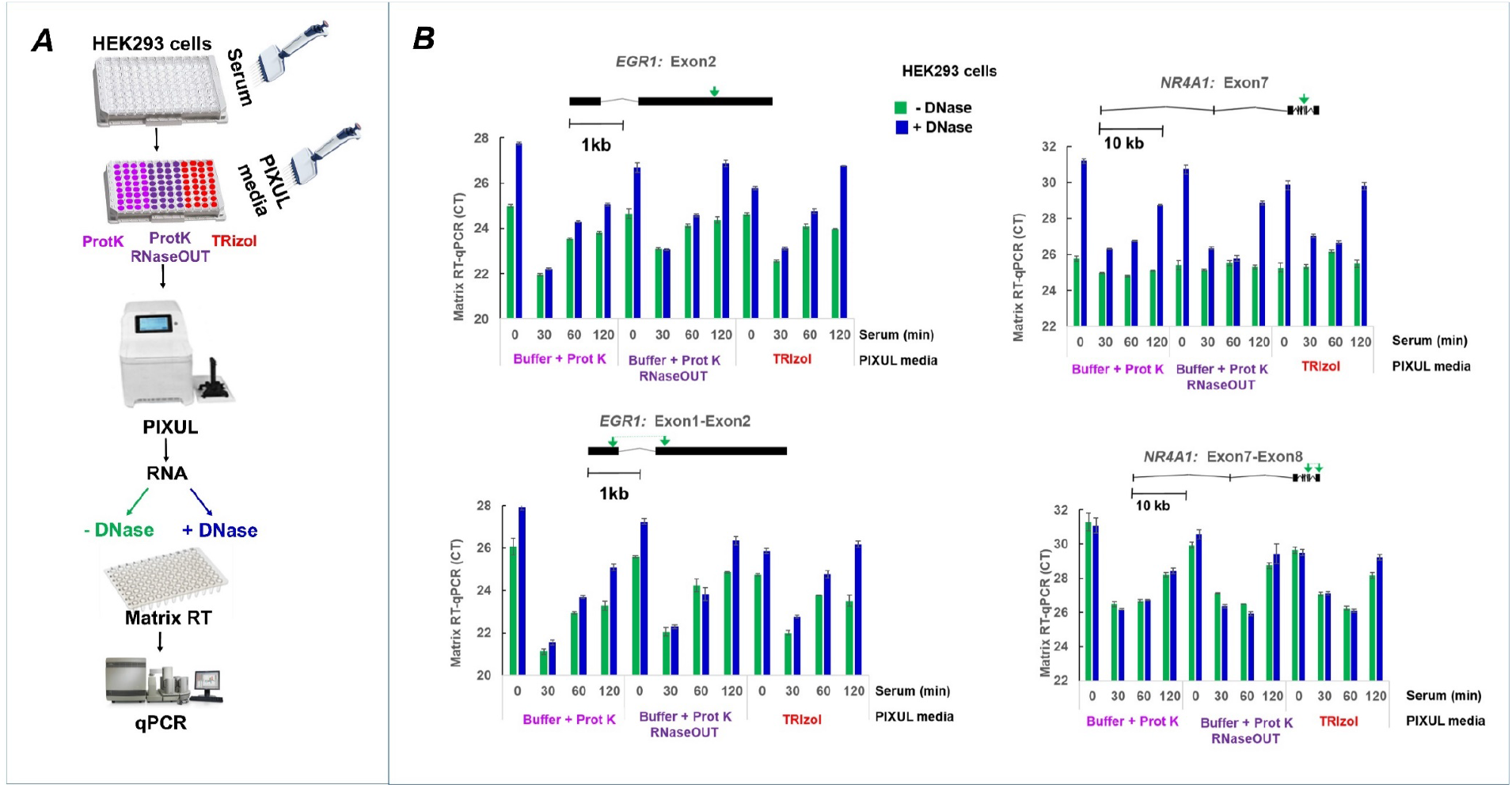
PIXUL-Matrix-RT-qPCR analysis with or without DNase of serum inducible genes in 96-well human HEK293 cultures. ***A*,** Serum-deprived HEK293 96-well cultures were treated with serum for 0, 30,60 and 120min. Media was aspirated and replaced with either elution buffer-proteinase K, with or without RNaseOUT, or TRIzol. Plates were treated in PIXUL, RNA was isolated and either treated (+DNase) or not (-DNase) with DNase I. RNA was used in Matrix RT-qPCR with indicated primers. ***B***, Cartoons above graphs show primers used in qPCR. Data are expressed as cycle threshold (CT).

Sepsis causes profound multiorgan changes in mRNA expression profiles, providing a system to test RNA isolation methods from organs using models of this syndrome (48–52). Given its substantial RNase levels, liver could be a challenging organ to efficiently extract RNA from while minimizing degradation. Next, we tested the above protocol using either proteinase K or TRIzol extraction from CryoCore samples taken from CryoTray-stored frozen livers of either LPS (endotoxin) or PBS (control) treated mice (**Fig.3A**). Isolated RNA was used in Matrix RT-qPCR using exon-spanning primers to the housekeeping *Actb*, and multiorgan sepsis-inducible *Ngal* (*Lcn2*) genes (48,51). The CTs for *Actb* were much higher (lower mRNA levels) with the proteinase K method compared to TRIzol (**Fig.3B**). Further, while there was LPS-induced substantial decrease in *Ngal* CTs in TRIzol-purified RNA, with the proteinase K method CTs were higher and there was only a small LPS-induced CT decrease (**Fig.3C**). The striking LPS-response differences between the two methods is further underscored when the *Ngal* mRNA levels are normalized to *Actb* (**Fig.3D**). These results suggest that the liver RNases degrade RNA even with the inclusion of RNaseOUT.

**Fig.3.**
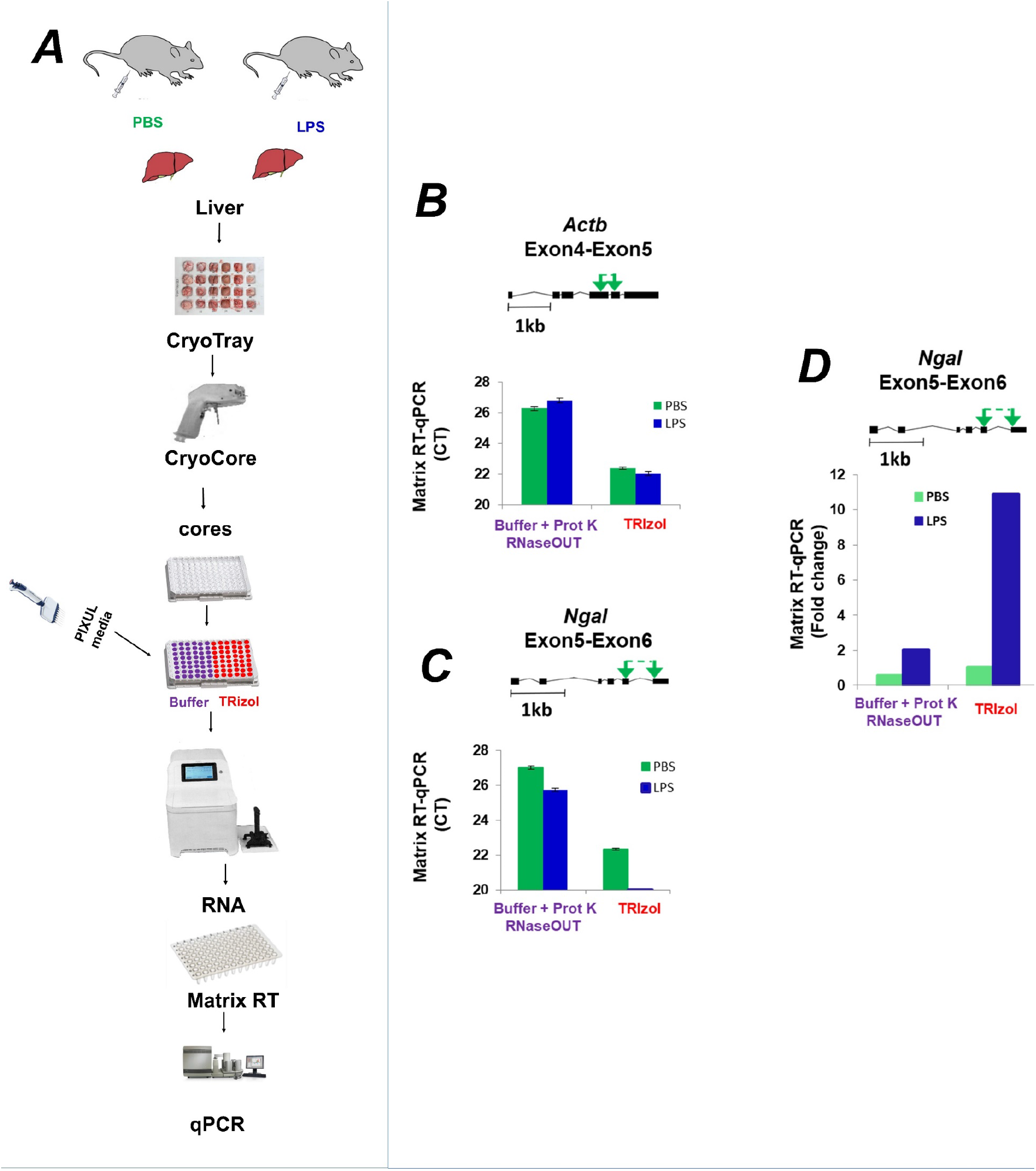
PIXUL-Matrix-RT-qPCR analysis of liver transcripts harvested from LPS, or PBS (control) treated mice and RNA isolated using either elution buffer-proteinase K or TRIzol from LPS. ***A*,** Mouse livers from LPS or PBS treated mice were frozen in CryoTray, sampled with CryoCore and jetted directly into wells of 96-well PIXUL plate. Samples were sonicated in either elution buffer-proteinase K or TRIzol. Isolated RNA was analyzed in Matrix-RT-qPCR. ***B-C*,** Cartoons above the graphs show mouse primers spanning either *Actb* (B) or *Ngal* (C) exons used in qPCR. Data are expressed as cycle threshold (CT). ***D, Ngal*** mRNA expressed as fold change relative to *Actb* mRNA.

To further explore the role of RNases, CryoCore mouse liver samples were ejected into wells with proteinase K buffer overlaying HEK293 cells from serum-treated cells. Plates were sonicated in PIXUL, RNA was isolated and *EGR1* mRNA levels were assessed in Matrix RT-qPCR using primers spanning Exon1-Exon2 region (**Fig.4A**). **Fig4B** shows that adding liver cores to the HEK293 cells increased CT (less RNA) values and no serum response was detected. These results provide evidence that the failure of the proteinase K buffer, which works well in HEK293 culture (**Figs. 2 and S3**), to isolate RNA from livers might in part reflect RNase degrading activity in this organ.

**Fig.4.**
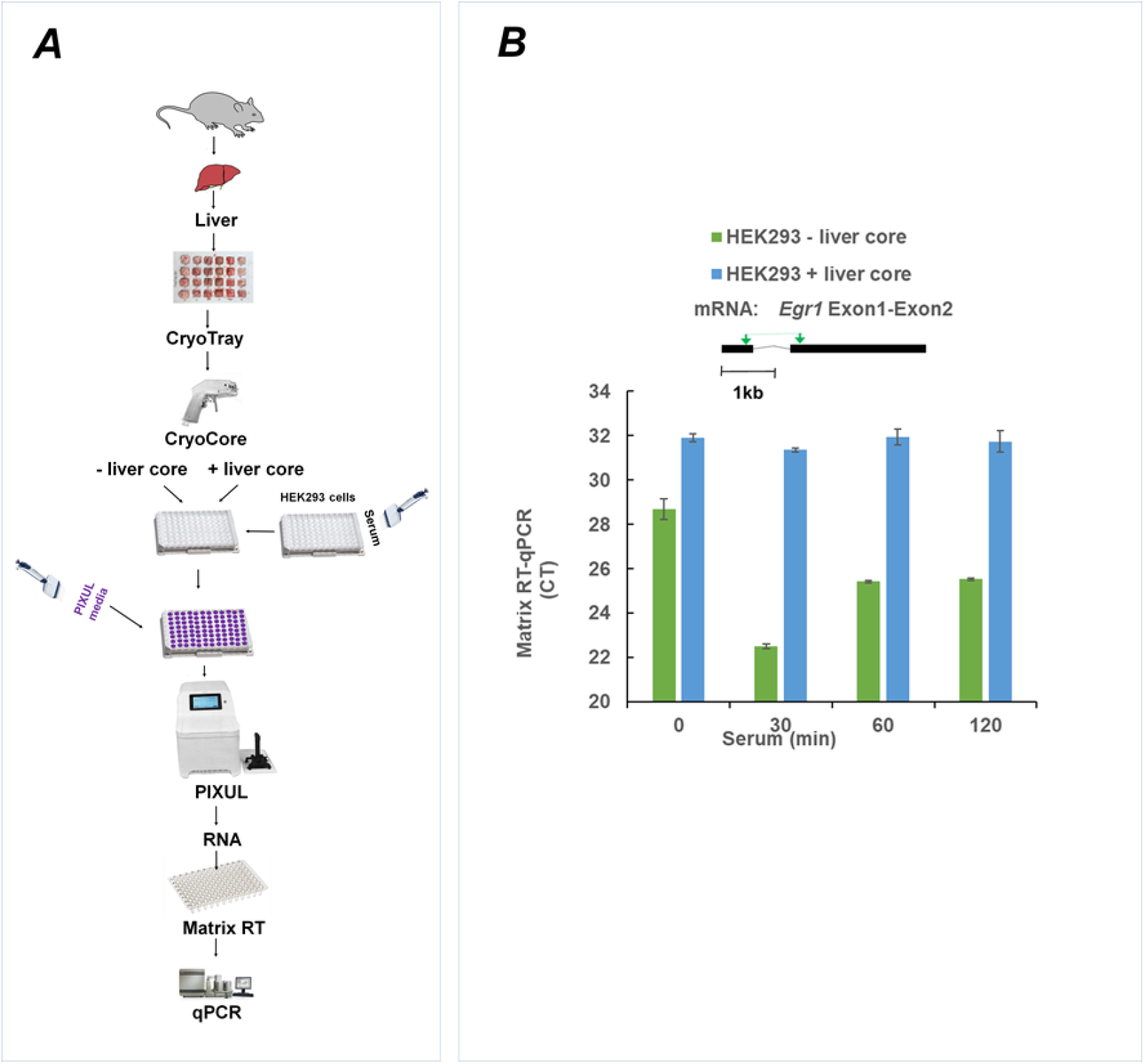
PIXUL-Matrix-RT-qPCR analysis of serum inducible *EGR1* gene in 96-well human HEK293 cultures co-sonicated with or without mouse liver cores. Serum-deprived HEK293 96-well cultures were treated with serum for 0, 30, 60 and 120min. Media was aspirated and replaced with elution buffer-proteinase K. CryoCore liver samples jetted or not to given wells, samples were sonicated in PIXUL, and RNA was isolated. RNA was used in Matrix RT-qPCR with indicated primers. Data is expressed as a cycle threshold (CT).

Using the above proteinase K buffer recipe we unsuccessfully tested other known inhibitors of RNases including polyvinylsulfonic acid (PVS) (53) and bentonite (54) to isolate high-quality RNA from mouse livers. To increase the activity of proteinase K and decrease the activity of RNAses we tested adding chaotropic reagents including SDS and guanidium hydrochloride. We tested a range of pH and concentrations of EDTA, DTT and SDS to maximize tissue RNase inhibition and preserve RNA integrity from mouse livers using the PIXUL proteinase K (PK) buffer protocols. SPRI are used to remove inhibitors. Given that SDS inhibits PCR and DNase activity we added SPRI beads step to isolate RNA and then treated the RNA prep with DNase I. Further, we added RNaseOUT to inhibit residual RNases during DNase treatment. The optimized PIXUL PK buffer and TRIzol protocols are shown in **Fig.5**. After PIXUL treatment TRIzol samples are moved under the hood to 500μl microcentrifuge tubes with an additional 100μl of TRizol (total = 200μl) and follow the TRIzol RNA extraction protocol followed by DNase I digestion. In PK buffer protocol samples are immediately incubated in the same polypropylene PIXUL plate at 95°C for 20 minutes, and then put on ice for 5 minutes. Cooled samples are centrifuged in a plate centrifuge for 10 minutes at 4,200 ×g and 4°C and then put back on ice. 60μl of clear supernatant (without any floating residue) from each well is carefully transferred via pipette to a new semi-skirted 96-well PCR plate on ice. Isolation of nucleic acids with 1.8x SPRI beads is done as per the Omega Bio-tek protocol (Norcross, GA) with a two-minute final SPRI bead drying time and final elution to 50μl of ultrapure ddH_2_O and then put samples on ice. Finally, DNase I and RNAse Out are added to the eluted RNA samples and DNase I digestion is done as per the manufacturer’s protocol.

**Fig.5.**
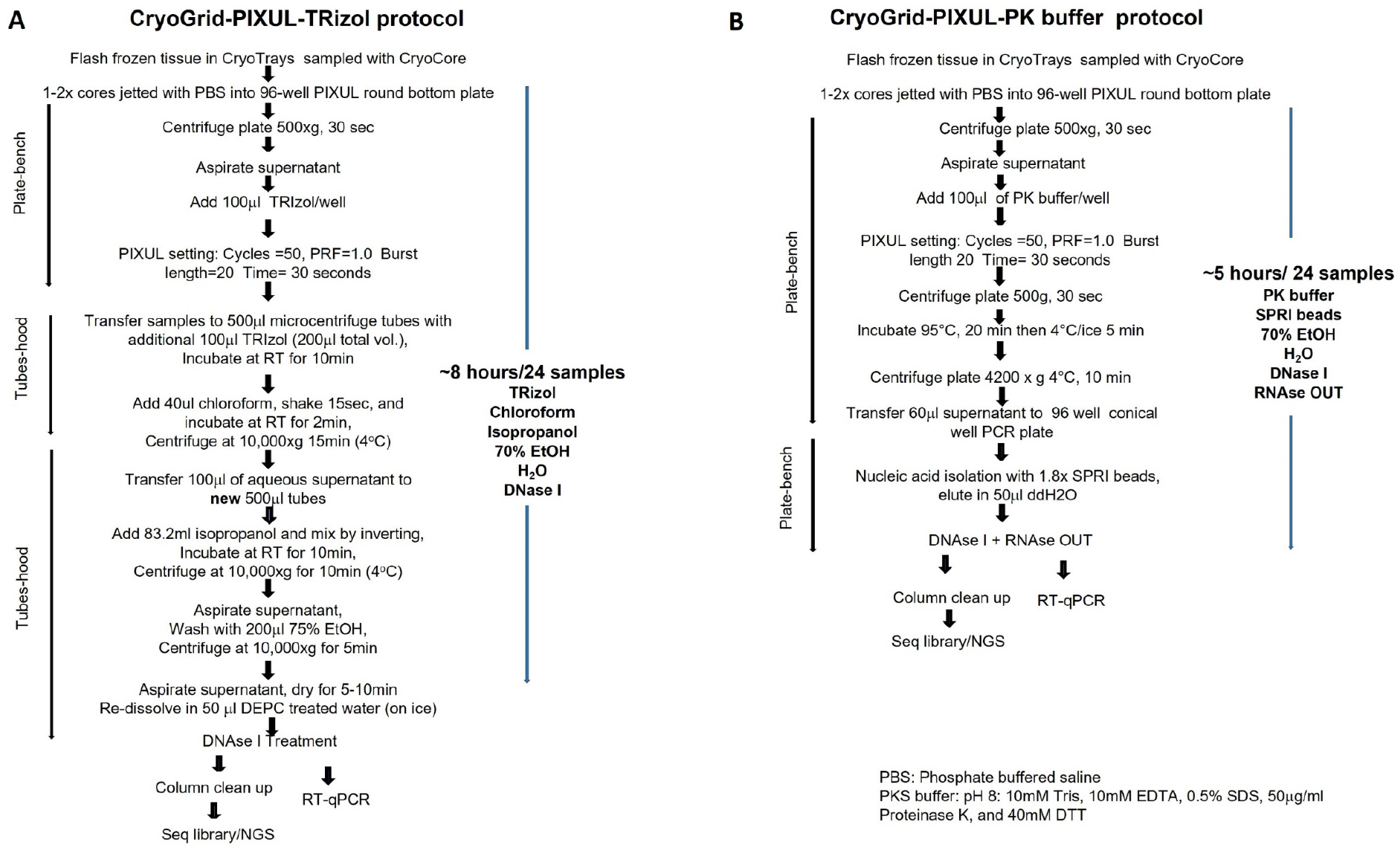
CryoGrid-PIXUL-RNA-qPCR/seq using TRIzol or PK buffer protocol for tissue RNA extraction. ***A***, The TRIzol protocol involves transfer from PIXUL 96-well plate to tubes and then another transfer to a new set of tubes. Besides TRIzol this protocol uses two other organic solvents, chloroform and isopropanol, steps that need to be done under the hood. It requires DNase I treatment to remove gDNA. This protocol takes approximately 8 hours to isolate RNA from 24 tissue samples. ***B***, The PK buffer protocol involves one transfer from PIXUL 96-well plate to a conical 96-well plate, does not use tubes or organic solvents and is done on an open bench. It requires SPRI beads, DNase I to remove gDNA and RNAseOUT to inhibit residual RNAses. This protocol takes approximately 5 hours to isolate RNA from 24 tissue samples.

### Comparing PIXUL-PK buffer and PIXUL-TRIzol RNA isolation methods in multiple mouse organs

We used organs from septic and control mice to compare RNA isolation using the PIXUL-PK buffer and PIXUL-TRIzol protocols. Mouse brains, hearts, kidneys, and livers were harvested 12hrs after either LPS (endotoxin) or PBS (control) intraperitoneal injection (IP) and frozen in CryoTrays. Organ cores were extracted with the CryoCore and jetted into two PIXUL plates, one for PK buffer and the other for TRIzol PIXUL treatment. RNA quality (absorbance 260/280 and 260/230) was estimated with the NanoDrop and RNA integrity number (RIN) was measured with the Agilent 4200 TapeStation system. Organs’ 280/260 (ranged 1.97-2.09) and 280/230 (ranged 1.80-2.35) absorbance ratios were nearly identical for the two methods (**Fig. S4A**). RIN, which is a qualitative – not quantitative – metric (Agilent Technical Support), is automatically derived using a proprietary algorithm originally developed for the Agilent Bioanalyzer. It assesses the entire electrophoretic trace of an RNA sample including the 18S and 28S area ratios (55). In the original Agilent study RIN was correlated with the quality of microarrays and RT-PCR data (55). In contrast to that original publication, there are microarray studies where only 1% of probes were correlated with RIN (56). Moreover, it has been shown that with selected library preparation protocols the number of mapped reads in RNA-seq demonstrated little or no correlation with RIN (57). Our studies have shown that RIN was lower with the PK buffer protocol compared to TRIzol for each organ as follows: brain 4.71+0.15 vs. 5.83+0.14 (p<0.01); heart 3.04+0.15 vs. 5.64+0.24 (p<0.01); kidney 2.84+0.15 vs. 4.13+0.15 (p<0.01); and liver 3.04+0.13 vs. 4.73+0.21 (p<0.01) (**Fig. S4B**). It is possible that RIN was lower for PK buffer compared to TRIzol method because it involves heating step (56) and/or the more fragmentation of RNA during PIXUL treatment. However, given that we used random hexamer priming, the PIXUL and PK buffer protocol have not reduced quality or RT-qPCR transcript analysis compared to conventional TRIzol RNA isolation method (below).

Matrix-RT-qPCR was used first to compare the PK buffer vs TRIzol protocol (**Fig.6A**) to assess the well-described organ-specific, sepsis-induced, and housekeeping mRNA expression patterns (1,48,51). Data are shown as both CTs and as a ratio to the *Rpl32* housekeeping mRNA (**Fig.6B**). *Syn1, Tnnt2, Fxyd2, Alb* show striking organ-specific signals for brain, heart, kidney and liver respectively. The housekeeping genes (*Actb* and *Rpl32*) did not show significant organ specificity. For organ-specific genes there was a moderate LPS-induced downregulation in the heart (*Tnnt2*) and kidney (*Fxyd2*) As illustrated by CTs, constitutive *Ngal* expression is much higher in the liver compared to the other organs. *Ngal (Lcn2*) expression was induced by LPS treatment in all four organs as illustrated by decreased CT and increased ratio to *Rpl32*. Elevated LPS endotoxin-induced *Ngal/Lcn2* expression in liver as well as inducible expression across multiple organs has also been shown in the mouse cecal ligation and puncture (CLP) sepsis model (48). The results obtained with the PIXUL-PK buffer were indistinguishable from those with the PIXUL-TRIzol method, suggesting that the two methods are equivalent when measuring expression of these genes by RT-qPCR (**Fig.6**).

**Fig.6.**
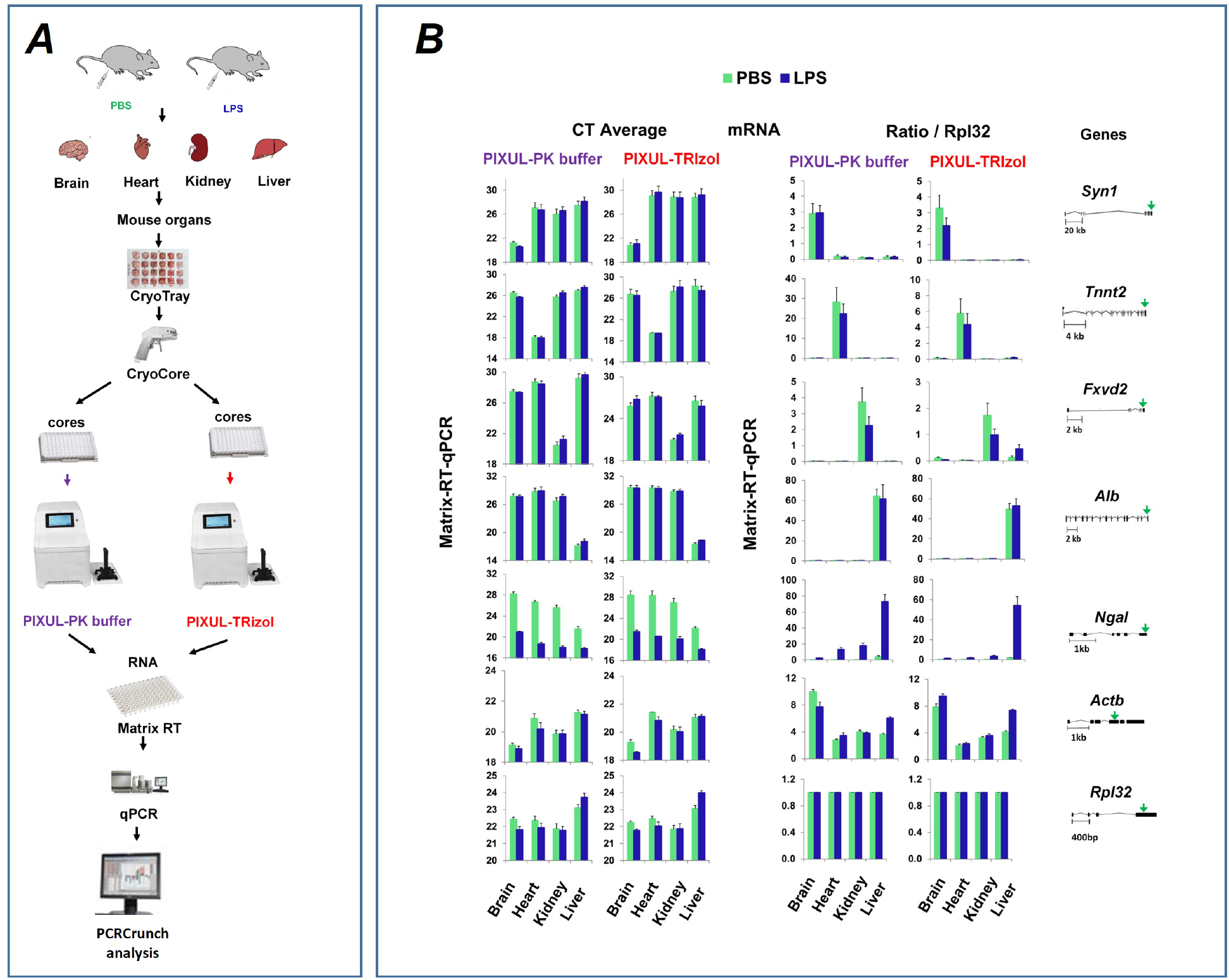
PIXUL-Matrix-RT-qPCR analysis of brain, heart, kidney, and liver transcripts from LPS (endotoxin) or PBS (control) treated mice and RNA isolated using either SDS buffer-proteinase K (PK buffer) or TRIzol. Mouse brains, hearts, kidneys, and livers from LPS or PBS treated mice were frozen in CryoGrid, CryoCore sampled, and then cores jetted directly into wells of 96-well PIXUL plate. Samples were sonicated in either SDS buffer-proteinase K (PK buffer) or TRIzol. Isolated RNA was analyzed in Matrix-RT-qPCR with primers to last exons of the indicated mouse genes. Data are expressed either as CTs (*left*) or normalized to *Rpl32* (*right*) housekeeping gene.

### RNA-seq analysis of mouse organs

RNA-seq provides a transcriptome-wide view at gene expression which we used next to compare the two tissue RNA extraction protocols. Zymo-Seq RiboFree Kit was used to prepare libraries which were then sequenced on Illumina NextSeq2000 (**Fig.7A**). All samples for both TRIzol and PK buffer methods of RNA isolation had mean quality per base Phred scores greater than 30 across all base pairs, equating to greater than 99.9% accuracy in all base calls (**Fig.S5A-B)**(24). RSeqQ gene body read coverage analysis (23) of RNA isolated with either TRIzol or PK buffer showed even distribution across the gene bodies (**Fig.S5C-D**) suggesting that coverage of transcripts using either isolation method is both uniform and nearly indistinguishable. Scatter plot analysis of RNA-seq raw count data showed excellent agreement between PK buffer and TRIzol RNA isolation methods, R = 0.984 - 0.992 (**Fig.S6**). Principal component analysis (PCA) of RNA-seq data showed clear organ-specific gene clusters, and overall PK buffer and TRIzol data also cluster together (**Fig.S7**). The IGV browser RNA-seq screenshots at the mRNA 3’-ends of the organ-specific *Synl, Tnnt2, Fxyd2, Alb*, the house-keeping, *Actb* and the LPS-responsive *Ngal* genes recapitulate the qPCR results (**Figs. 6B&7B**). As in the case of the RT-qPCR data (**Fig.6B**) the RNA-seq results for these genes are indistinguishable for the two PIXUL RNA isolation methods (**Fig.7B**).

**Fig.7.**
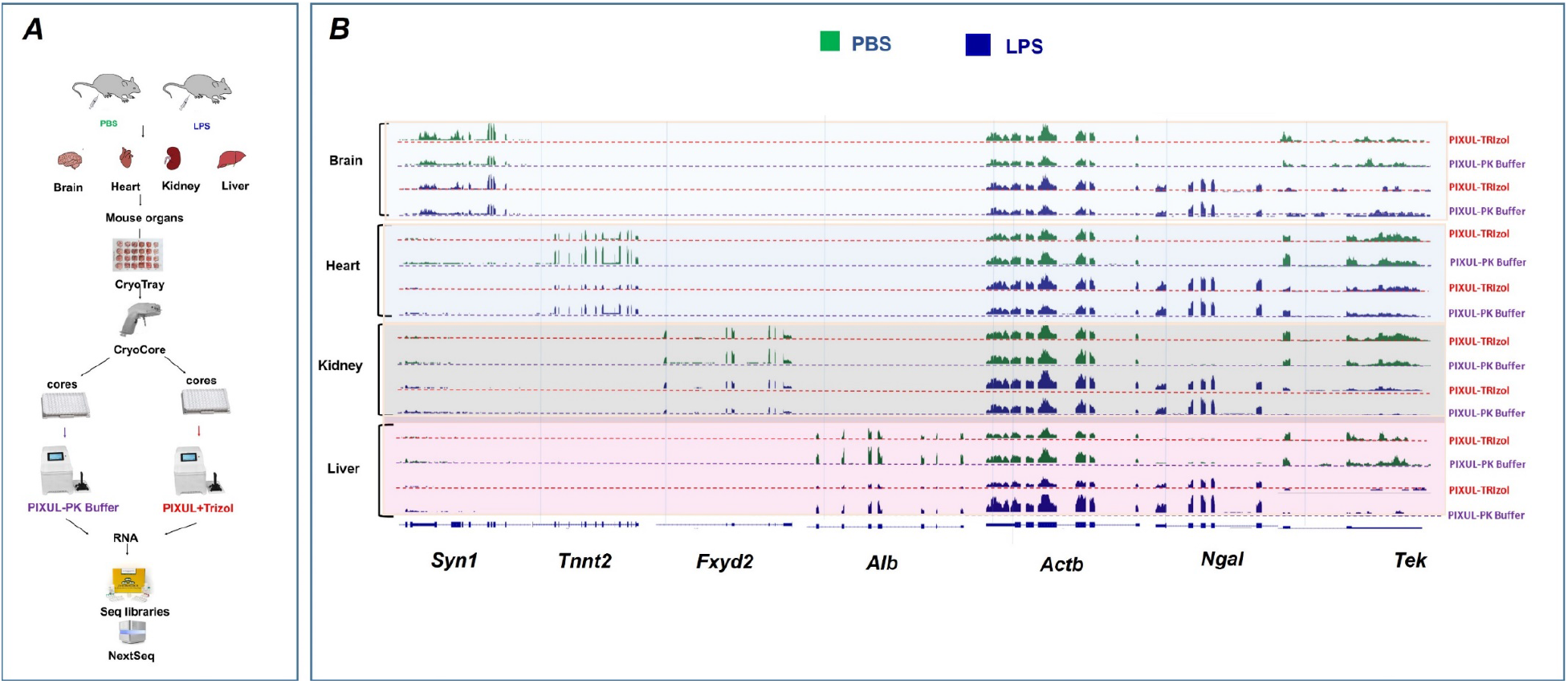
CryoGrid-PIXUL-RNA-seq analysis of brain, heart, kidney, and liver transcripts from LPS or PBS treated mice and isolated using either PK buffer or TRIzol. ***A*,** Mouse brains, hearts, kidneys, and livers from LPS or PBS treated mice were frozen in CryoGrid, CryoCore sampled and then jetted directly into wells of 96-well PIXUL plate. Samples were sonicated in either PK buffer or TRIzol. Sequencing libraries were generated using Zymo-seq RiboFree kit and sequenced on NextSeq2000 employing a dual-index, paired-end, 61 base read length (PE61). ***B*,** IGV mRNA tracts for organs specific at 3’-ends of organ-specific (brain: *Syn1*, heart: *Tnnt2*, kidney: *Fxyd2* and liver: *Alb*), house-keeping (*Actb*), sepsis-inducible (*Ngal*(*Lcn2*)) and sepsis-down-regulated (*Tek*) genes. Data shown represent one RNA-seq analysis out of two independent experiments.

We carried out differential gene expression (DGE) analysis with ‘EdgeR’ as another way to compare the two RNA isolation methods. Mean difference (MD) plots comparing PK buffer vs. TRIzol RNA isolation methods show very few statistically significant differences in differentially expressed transcripts between the two methods (**Fig.S8A**). In contrast to this comparison, MD plots of transcripts isolated with either PK buffer or TRIzol method illustrate large numbers of statistically significant LPS-induced or repressed genes (LPS vs. PBS comparisons) in the heart, kidney, and the liver but there were much fewer LPS-responsive genes in the brain (**Fig.S8B**). Numerical values are shown in **Fig. S9**. With both methods, after filtering lowly-expressed genes, there were more than 21,000 transcripts detected in each one of the four organs (**Fig. S9A-C**). The number of transcripts for each organ differentially expressed in TRIzol but not PK buffer RNA preps were at most 6 (heart) while those differentially expressed in PK buffer but not TRIzol were at most 4 (kidney). The number of LPS-(upregulated//downregulated) transcripts with PK buffer vs. TRIzol were as follows: brain 119//7 vs. 237//42; heart 820//468 vs. 1020//599; kidney 1594//1136 vs. 1064//610; liver 1593//1622 vs. 1994//2246 (**Fig.S9D-F**). Thus, the percent of genes upregulated//downregulated in response to LPS was as low as <1%//1% in the brain but as high as 10%//10% in the liver. The low endotoxin response in the brain is not unexpected given the blood-brain barrier. In each organ there were LPS-induced DGEs seen using TRIzol but not PK buffer and vice versa. But in each case (except for the brain) there were substantially more LPS DGEs that were shared by the two methods than not (**Fig.8A**). Pearson’s correlation coefficient for DGEs in organs between the RNA isolation methods were as follows: brain 0.777; heart 0.922, kidney 0.846 and liver 0.828 (**Fig.8B**).

**Fig.8.**
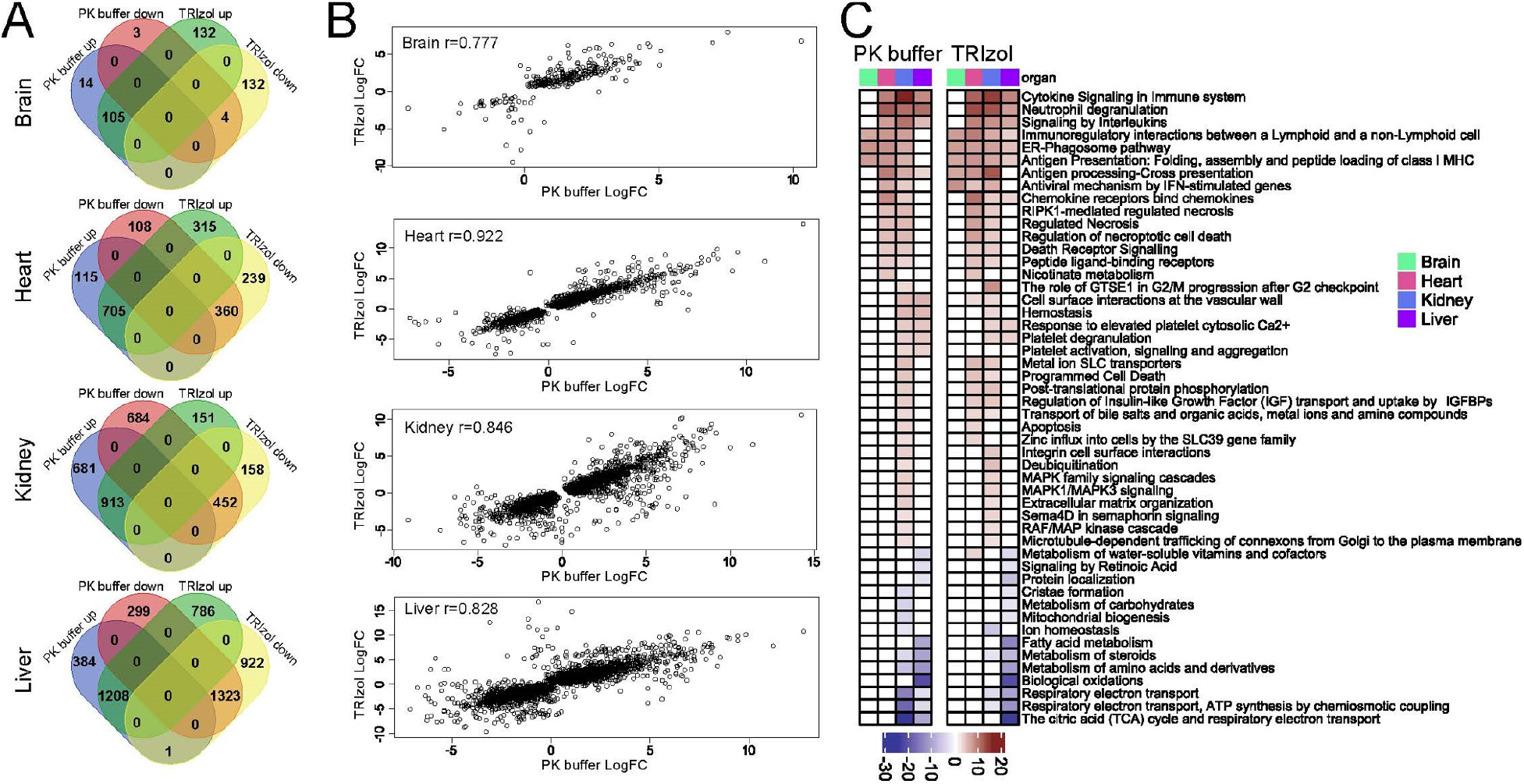
Differential gene expression (DEG) and Pathway analysis. ***A*,** Venn diagrams depicting the overlapping numbers of up-and downregulated DEGs (adj. p-value < 0.05) for mouse organs following LPS endotoxin challenge between the TRIzol and PK buffer RNA isolation methods. ***B*,** Pearson’s correlation coefficient for DEGs in organs between the two RNA isolation methods. ***C***, Significantly altered Reactome pathways (adj. p-value < 0.05) across the organs following LPS endotoxin IP injection for the TRizol and PK buffer RNA isolation methods. The Reactome pathways with a differential gene ratio higher than 10% are shown. The p-value is on the PHRED scale and is reversed for down-regulated pathways.

Reactome PK buffer and TRIzol RNA-seq data analysis of LPS-induced DEGs for each organ identified nearly identical pathways well described in models of sepsis (48,50,58). This included such pathways as cytokine signaling in the immune system, neutrophil degranulation, signaling by interleukins, platelets degranulation and several others (**Fig.8C**). Altogether these analyses using RNA-seq data demonstrate that the two CryoGrid-PIXUL based methods of RNA extractions yield similar results.

### Limitations

The mouse heart, the smallest organ (<150mg) used in this study, can be sampled several times with the CryoCore (**Fig.S2**). Still, there are situations where the available tissue fragments are smaller than the mouse heart (e,g, clinical samples). Here, to be effective, miniature trephine gauges will be needed for multisampling very small tissue fragments in research and clinical settings (e.g. for genomics, epigenetics, transcriptomics, proteomics and histology). The current CryoTrays hold 24 tissue fragments. For small tissue samples, CryoTrays with 48 or 96-wells would further save cryostorage space.

Human operators are prone to error (59). Although using the iPad display largely mitigates human error using the 24-well CryoTray, with 48 or 96 samples on the CryoTray the introduction of computer vision and/or automation would provide an error-free sampling method. In this regard, we suggest that the application of the CryoGrid-PIXUL-Matrix-RNA-qPCR system to assays of organ-specific genes (**Fig. 6**) provides a convenient way to develop and test computer vision technology-assisted high throughout tissue sampling (60,61).

Although slow and taxing when using multiple samples, TRIzol is considered the “gold standard” method for RNA isolation (16) which we used as the benchmark in this study. The RIN was higher with TRIzol compared to PK buffer (**Fig.S4B**). Nonetheless, RNA-qPCR and RNA-seq results obtained with the PK buffer were similar to TRIzol (**Figs.6–7 and S5–S9**). But the PK buffer method has the advantage of being faster, biosafe, less labor-intensive and – given that it uses microplates – more suitable for automation. Still, there could be RNA species (e.g. small RNAs) where TRIzol performs better than the PK buffer protocol, or vice versa. More detailed comparative analysis is needed to find putative RNA species preferentially lost in one protocol versus the other protocol.

## CONCLUSIONS

After protein and DNA, RNA is the most studied biomolecule (PubMed, Google Scholar). RNA is increasingly being used as a clinical diagnostic analyte (62). Understanding RNA biology in tissues is critical in health and disease as well as therapeutic interventions (e.g., small molecules (63) and the rapidly emerging field of epigenome editing (64)). Freezing of tissues is widely used in research and clinical settings because it provides a practical way to preserve biospecimens for analysis. And yet, sampling of frozen tissues remains tedious. We developed a user-friendly cryostorage method and a hand-held sampling tool to interrogate tissues, CryoGrid system (**Figs. 1, S1–S2**), which we integrated with PIXUL sonicator for high throughput tissue RNA extraction. The CryoGrid-PIXUL-RNA workflow offers several advantages over existing platforms as follows.

i. It is a user-friendly method to freeze, track and sample tissues.
ii. One CryoCore 1-2 mm^3^ (1-2mg) tissue sample is sufficient for RNA-seq, thus allowing the same frozen tissues (e.g. as small as mouse heart) to be sampled multiple times for other analyses as well as histology.
iii. After sampling, and unlike commercially available RNA extraction platforms, the separate preceding tissue homogenization step is not needed.
iv. Virtually every disease and therapeutic agents’ effects are systemic conditions. The CryoGrid-PIXUL platform is well suited for parallel processing of multiple organs from model systems and for testing multi-organ effects in pre-clinical studies (52,64,65).
v. Ability to process dozens of samples at the same time mitigates batch effects (66).
vi. Transcriptomic, epigenetic, and proteomic analytes preparation can be done in parallel on the same PIXUL plate.
vii. The PK buffer method provides a way to isolate RNA without the use of hazardous solvents and as such can be done on the bench – a safety hood is not needed.
viii. Unlike TRIzol protocol, which uses tubes, the PK buffer RNA extraction procedure is done in 96-well plates, making it adaptable for automation.
ix. Has the potential to be used for tissue RNA extraction for single-cell (scRNA)-seq.
x. Cells or organoids grown in 96-well plates can be used directly to extract RNA in PIXUL without sample transfer (1).
xi. Using this system will expand the role of RNA-seq in pre-clinical studies and clinical settings (62).
xii. The CryoGrid system and the TRIzol-free RNA isolation methods can be easily used with a variety of sample preparation platforms.

## Data availability

Sequence data was deposited in Gene Expression Omibus database under entry GSE199598.

## Acknowledgments

This work was supported by NIH R42HG010855 and U01CA246503 (KB). We thank Dr. Mary Regier, UW ISCRM Genomics Core, for assistance with bioinformatics.

## Author Contributions

KB, conceived the project, designed the experiments, engineered the CryoGrid system, analyzed the data and wrote the paper. SAS, designed and carried out the RNA experiments, ran and analyzed RNA-seq and wrote the paper. SJS, engineered and built the CryoGrid system. DM, carried out experiments, contributed to CryoGrid design and edited the paper. AL, designed and carried out RNA experiments. OD, assisted in experimental design and edited the paper. WAA and DA did the mice experiments and harvested the mouse organs. TM, provided ultrasound expertise. MM and MK carried out RNA-seq pathway analysis.

## The authors declare the following competing financial interest(s)

KB and TM are co-founders of Matchstick Technologies, Inc. KB and TM are co-inventors of PIXUL (US Patent 10809166). KB, SJS and DM are co-inventors of CryoGrid components (patents applications 20210325280, 20210386056). KB and DM are co-inventors of the PlateHandle (patent pending). The above technologies are co-owned and/or have been licensed to Matchstick Technologies, Inc from the University of Washington.

## Inclusion and Diversity

One or more of the authors of this paper self-identifies as an underrepresented ethnic minority in science. One or more of the authors of this paper self-identifies as a member of the LGBTQ+ community. One or more of the authors of this paper self-identifies as living with a disability. One or more of the authors of this paper received support from a program designed to increase minority representation in science.

## SUPPLEMENTARY FIGURES

**Fig.S1.**
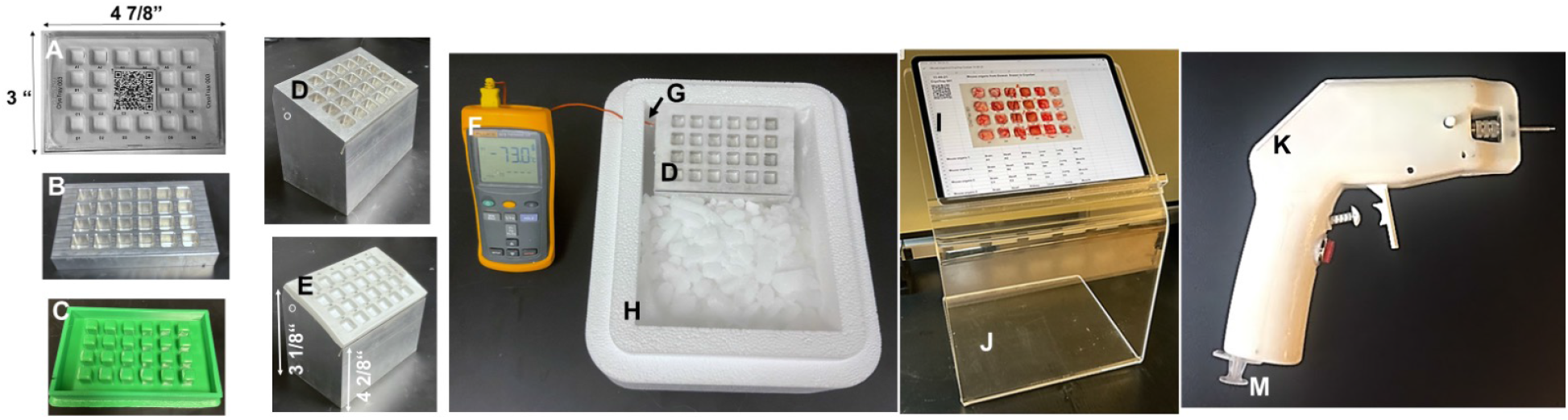
CryoGrid system components. 24-well CryoTray dimensions of a standard (4 7/8” x 3 1/8 “) plate (*A*) are casted in-house by placing polystyrene sheets (0.4mm) on heated (135°C) 24-cuboid sockets machined aluminum block (*B*) and then overlaying and pressing the sheet with a matching 3D printed 24-cuboid plugs plastic stamp (*C*). The CryoTray dimensions match an off-the-shelf clear polystyrene microplate lid (Cytivia #7704-1001). Online generated QR code sticker is secured on the inside of the lid and linked to Google Drive applications. CryoBlock is a 24-pockets machined aluminum block with a tilted top surface (30°) to improve ergonomics (*D*) which serves as a CryoTray receptacle (*E*) for freezing tissues and providing easy and open access for coring tissues. Fluke thermocouple thermometer (*F*) with a thermocouple probe (*G*) inserted into an aperture (shown in white circle) on the side of the CryoBlock (*D*) monitors temperature (<−70°C). CryoBlock is chilled in an off-the-shelf 11”x11”x9” Styrofoam box (CryoBox) containing dry ice pellets (*H*). iPad tablet screen (*I*) on an acrylic stand (*J*) displays online Google Sheet with a layout of frozen tissues and metadata of a file uploaded to the tablet by scanning CryoTray QR code to guide tissue sampling with the CryoCore (*K*). CryoCore is a miniature battery-powered tissue coring drill with a trephine burr. Trephine is composed of stainless steel tubing shaft (2mm x 30mm) and a smaller diameter tip (1mm x 1-2mm) which has axially oriented cutting teeth around the circumference. The length of the tip controls the tissue coring depth. The posterior (back) trephine end is mounted in the chuck of the rotary motor. A push-button switch (*red*) activates the motor allowing one to drill into the tissue fragment and then extract a core. Trephine has two holes with axes oriented perpendicular to shaft. A clamping mechanism uses two segments of silicone tubing with semicircular notches, which are equal in radius to the shank of the miniature coring drill, at their ends to close around the shaft holes, creating a seal. The clamp is actuated by a cable in a manner similar to a bicycle caliper brake. A pump mechanism utilizes a plunger and cylinder and two check valves to both draw buffer (e.g. PBS) from a 5ml refillable syringe reservoir (*M*) and to generate a fluid jet through the shaft holes to eject the extracted tissue core (67,68).

**Fig.S2.**
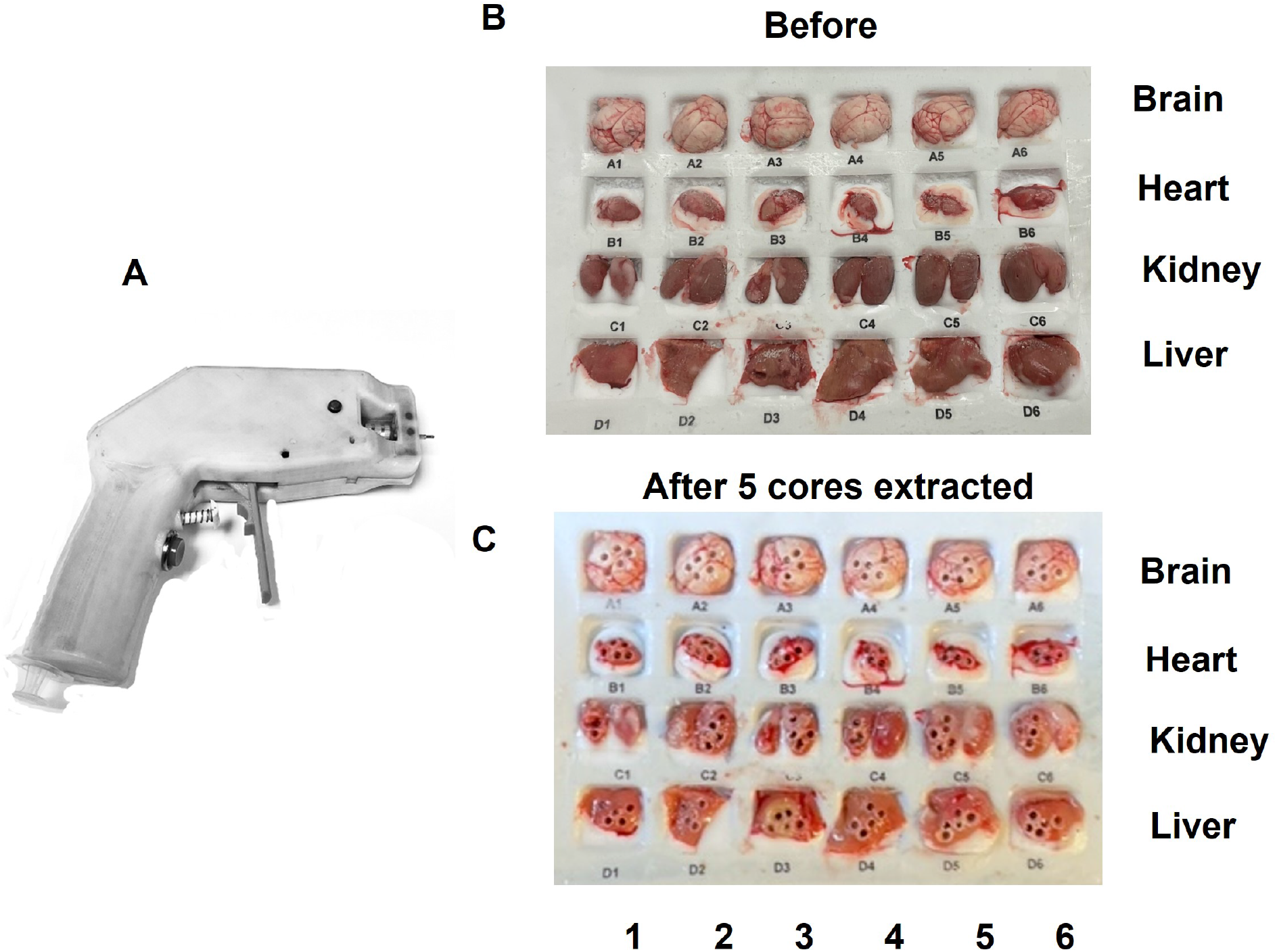
CryoTray with tissues before and after multiple CryoCore sampling. ***A***, CryoCore was used to sample mouse organs. ***B-C***, Frozen mouse organs in CryoTray were photographed before and after multiple CryoCore sampling, illustrating the size (~1-2 mm^3^) and the number of cores that can be extracted from an organ as small as the mouse heart (<150mg).

**Fig.S3.**
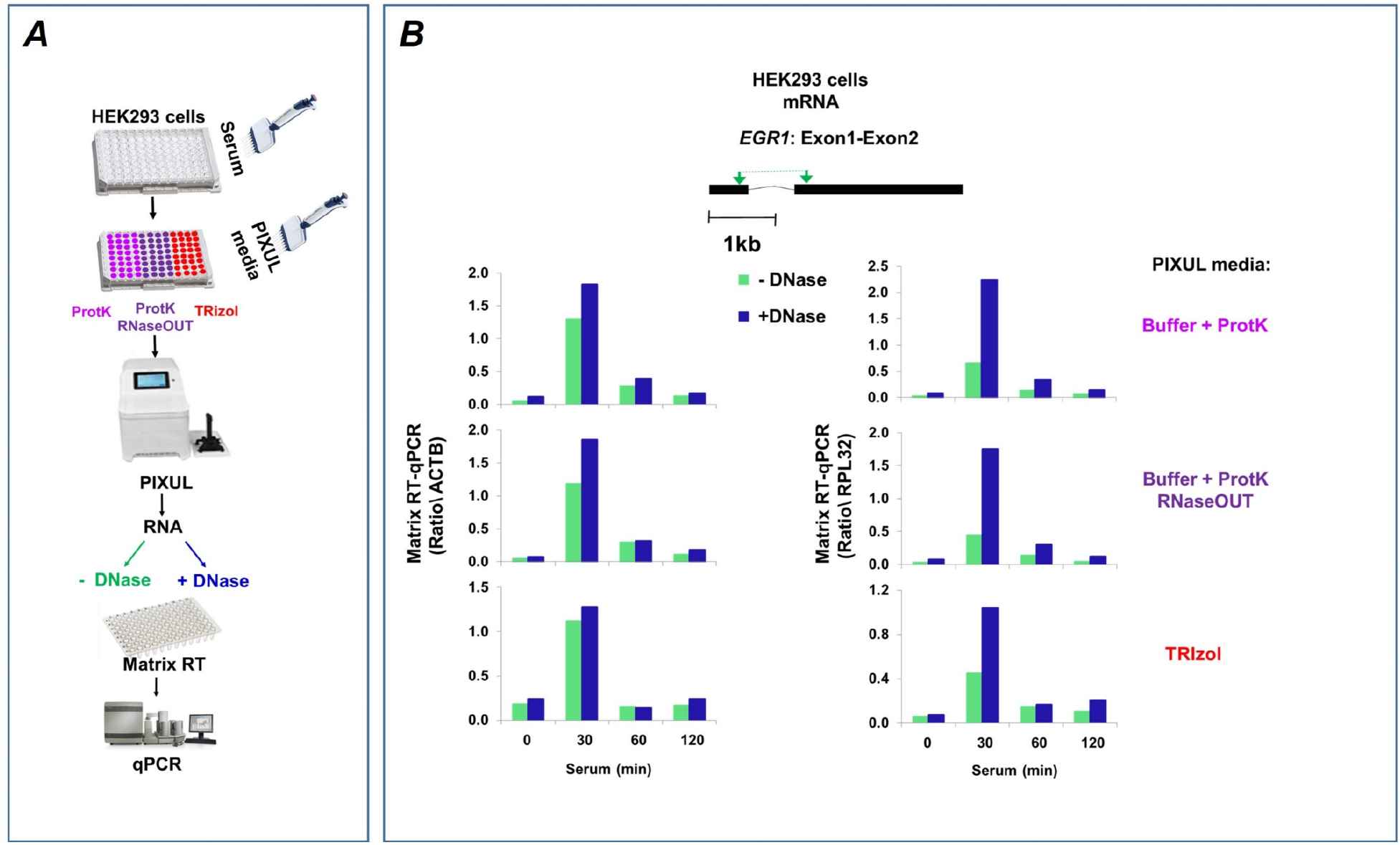
PIXUL-Matrix-RT-qPCR analysis with or without DNase of serum inducible *EGR1* in 96-well human HEK293 cultures. ***A*,** Serum-deprived HEK293 96-well cultures were treated with serum for 0, 30,60 and 120min. Culture media was aspirated and replaced with either elution buffer-proteinase K, with or without RNaseOUT, or TRIzol. Plates were treated in PIXUL, RNA was isolated and either treated (+DNase) or not (-DNase) with DNase I. RNA was used in Matrix RT-qPCR. ***B*,** Human *EGR1* primer spanning Exon1-Exon2 used is shown. *EGR1* mRNA data are expressed as a ratio to *ACTB (left*) and *RPL32 (right*) genes.

**Fig.S4.**
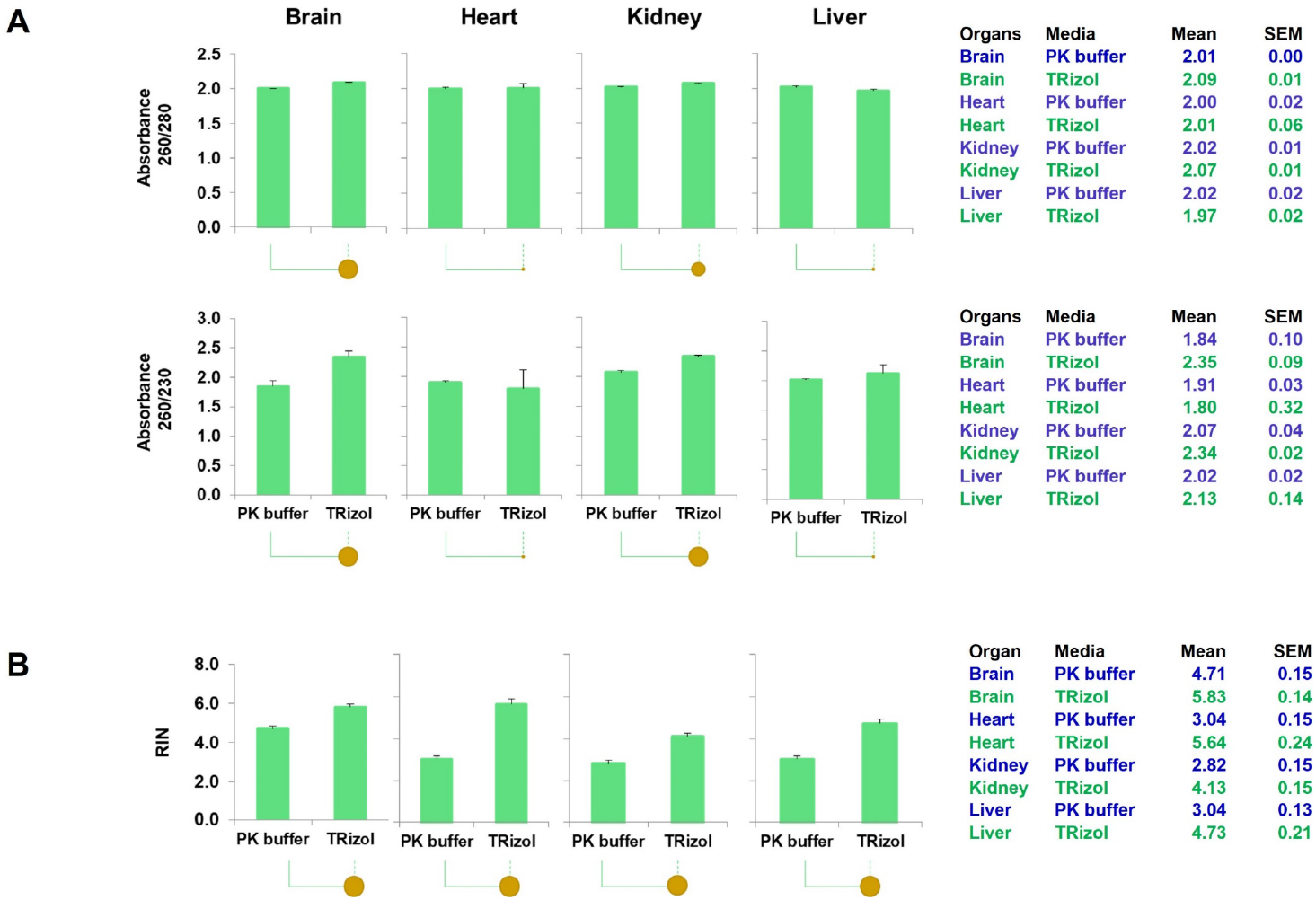
Quality of RNA isolated with PIXUL using PK buffer vs. TRIzol from mouse brain, heart, kidney, and liver. Frozen mouse organs in CryoTrays were sampled with the CryoCore and RNA was isolated using PIXUL with either PK buffer or TRIzol. ***A*,** NanoDrop absorbance is shown as 260/280 (*upper row*) and 260/230 (*lower row*). ***B*,** High Sensitivity RNA ScreenTape^®^ was used to assess the RNA integrity numbers (RIN) (55). Data are represented as mean + SEM, n=2 for each organ. Statistical differences between PK buffer and TRIzol (*P*-value) are shown by the size of the solid circle under the x-axis; P<0.05 by a medium circle, P<0.01 by a large circle, the tiny circle indicating the differences are not statistically significant (21).

**Fig.S5.**
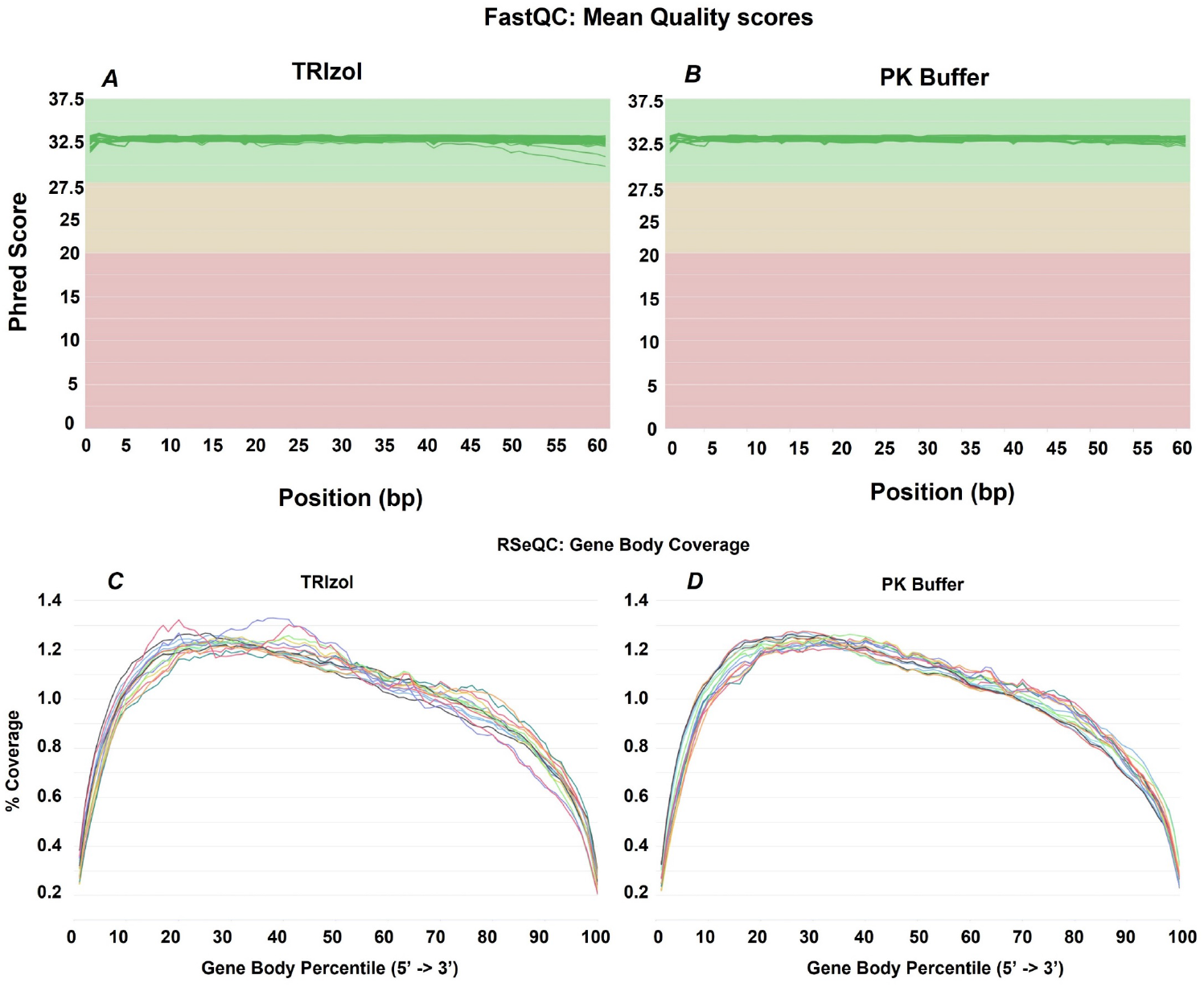
Sequencing quality. ***A-B*,** MultiQC compilation of FastQC ‘Per Base Sequence Quality’ scoring of RNA-seq quality for TRIzol (A) isolated samples and PK buffer (B) isolated samples. Each chart showing Phred score on the Y-axis and base pair position on the X-axis. ***C-D***, RSeqQC ‘Gene Body Coverage’ of read coverage percentage of TRIzol (C) and PK Buffer (D) isolated samples’ along the length of gene bodies that were normalized to each other (X axis, *Gene Body Percentile*)

**Fig.S6.**
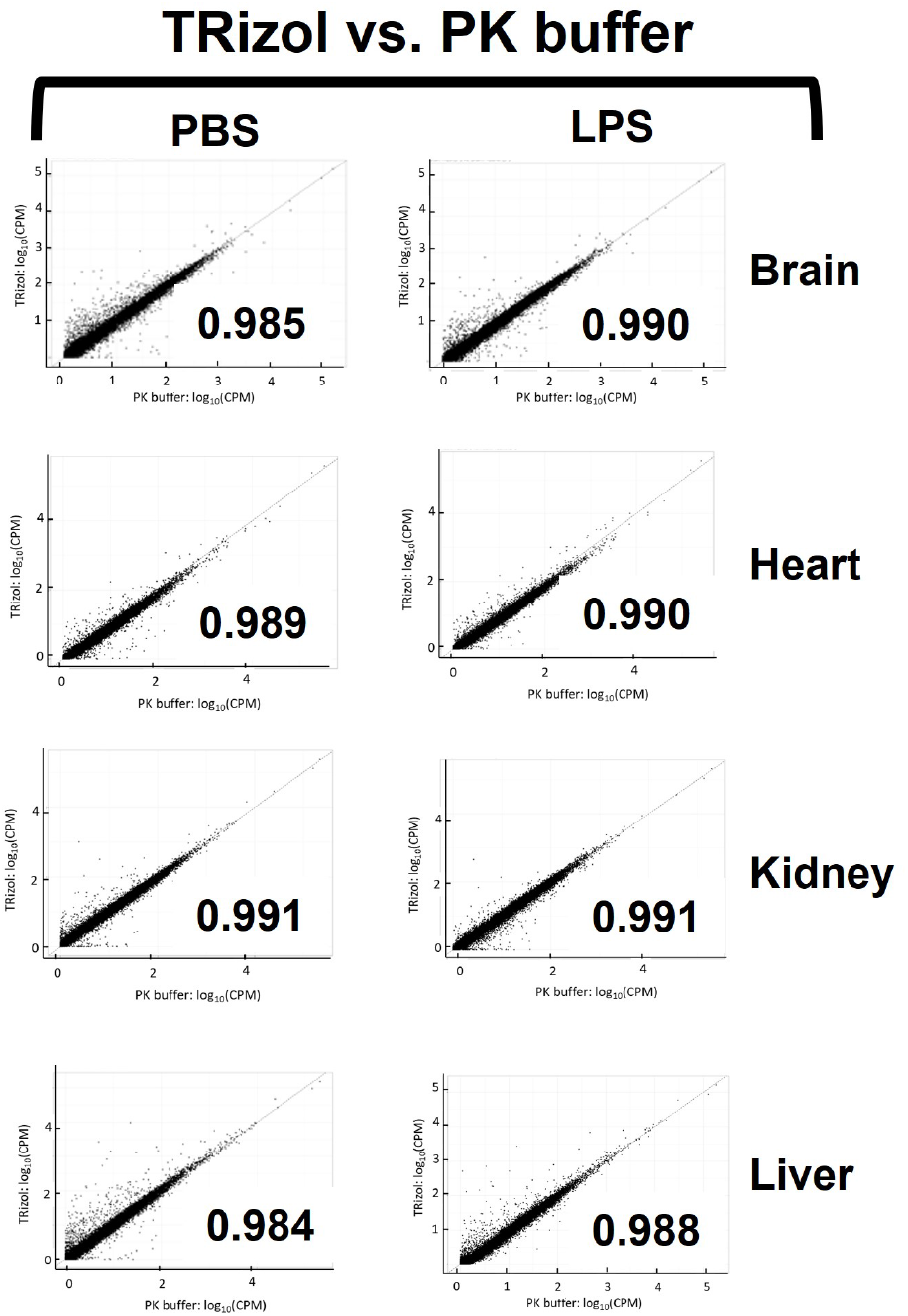
Scatter plots of correlation analysis of PK buffer and TRIzol of brain, heart, kidney and liver of RNA-seq raw-counts data from septic (LPS IP injected) and control (PBS IP injected) mice. Results are shown as log_10_ counts per million reads mapped (CPM), *x*-axis PK buffer and *y*-axis TRIzol. Number in the body of the graphs show correlation coefficients. Each dot represents a mean of two mice experiments.

**Fig.S7.**
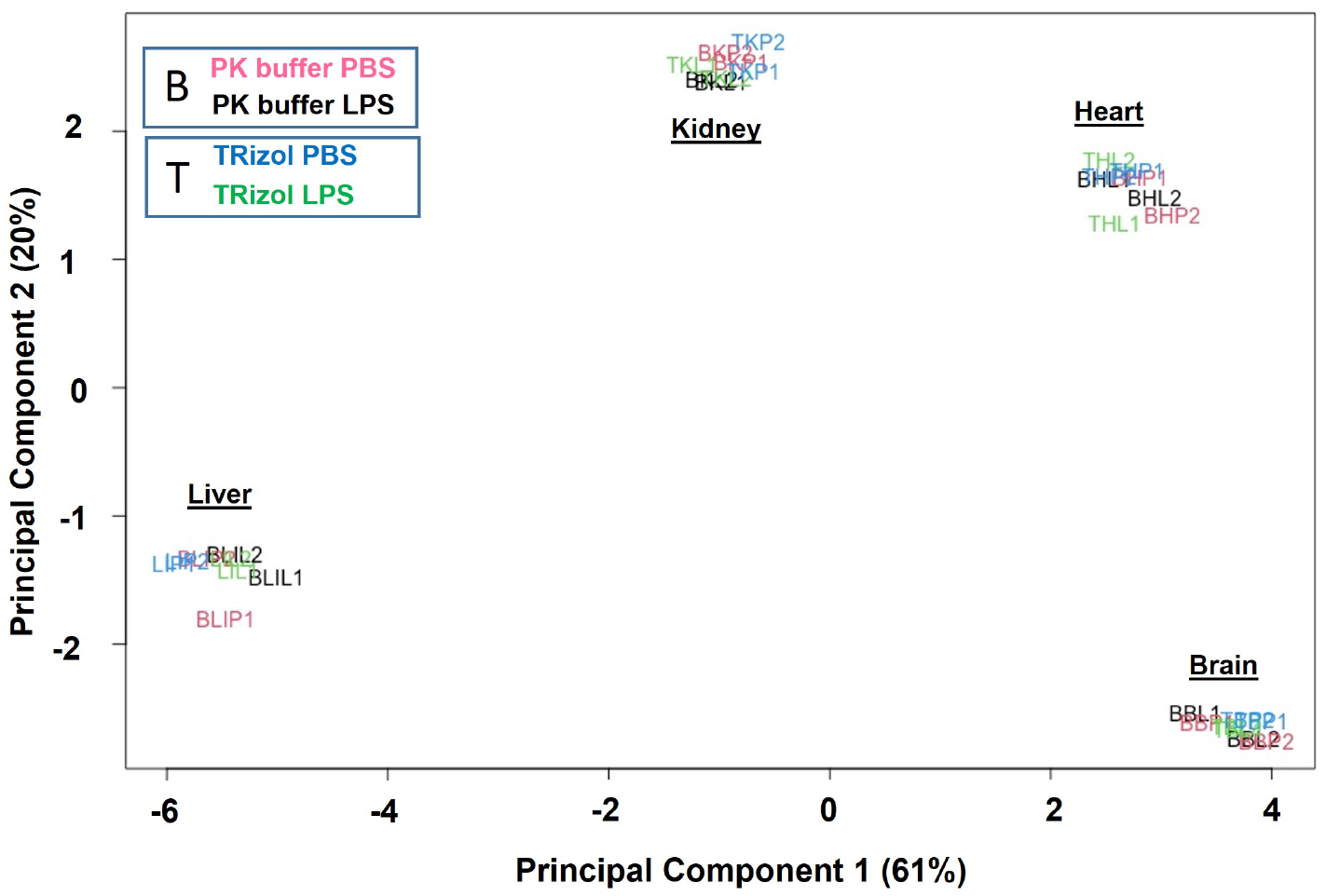
Principal component analysis (PCA) of brain, heart, kidney, and liver RNA-seq using RNA isolated with the PIXUL-PK buffer and PIXUL-TRIzol method from LPS and PBS IP injected mice. Data show two replicates for each organ for LPS and PBS IP-treated mice.

**Fig.S8.**
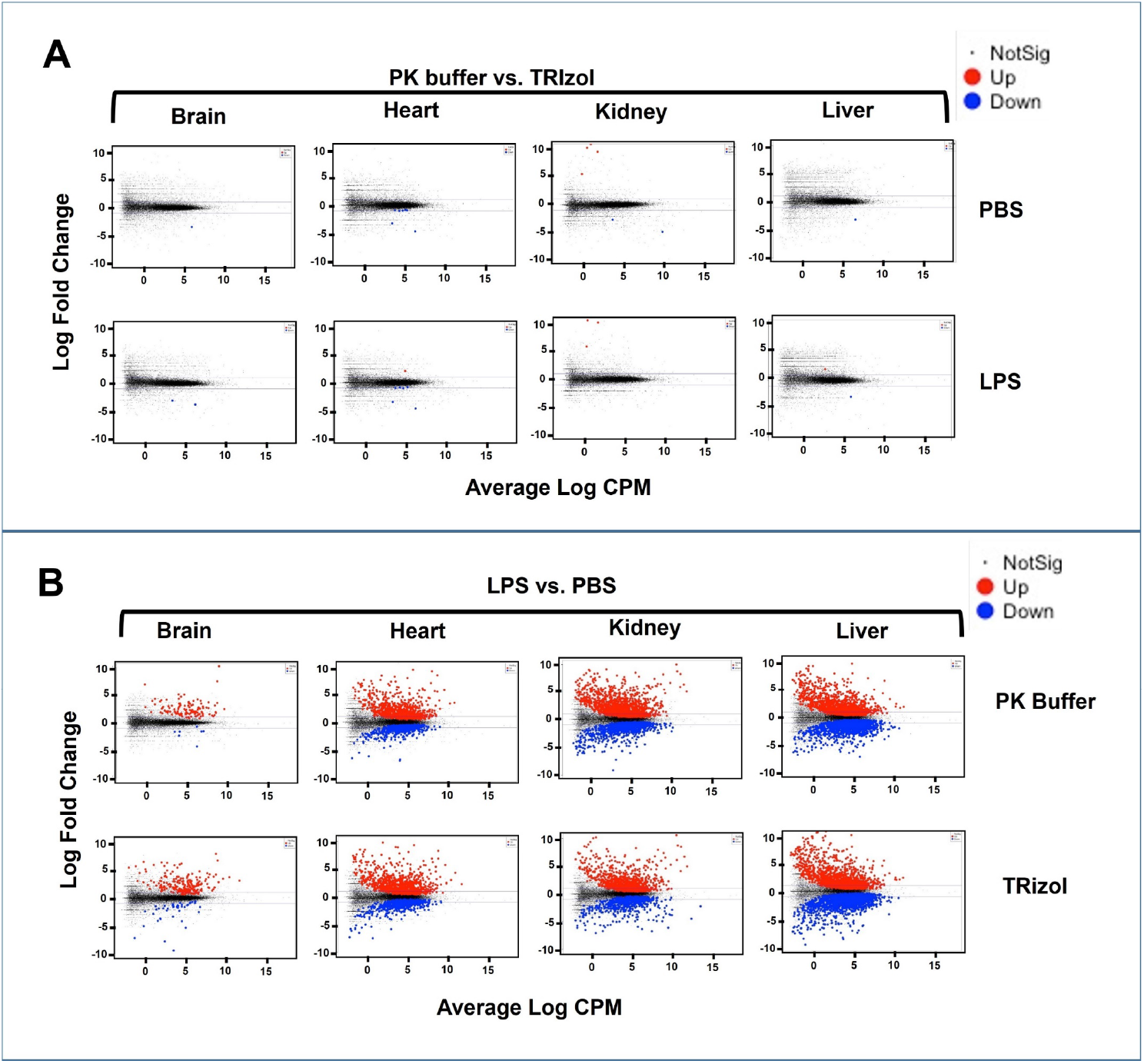
‘EdgeR’ Differential gene expression (DGE) shown with mean-difference (MD) plots. MD plot shows log fold change, y-axis and average log of counts per million (CPM), x-axis. Red and blue dots indicate transcripts that are statistically increased or decreased, respectively. Black dots represent transcripts that were not significantly differentially expressed. ***A***, transcripts extracted with PK buffer compared to TRIzol from brain, heart, kidney and liver from PBS-(upper panel) and LPS-treated mice. ***B***, comparison of brain, heart, kidney, and liver transcripts from LPS-treated compared to control (PBS) treated mice, *upper panel* PK buffer and *lower panel* TRIzol extracted RNA, respectively. y-axis log fold change; x-axis average log CPM.

**Fig.S9.**
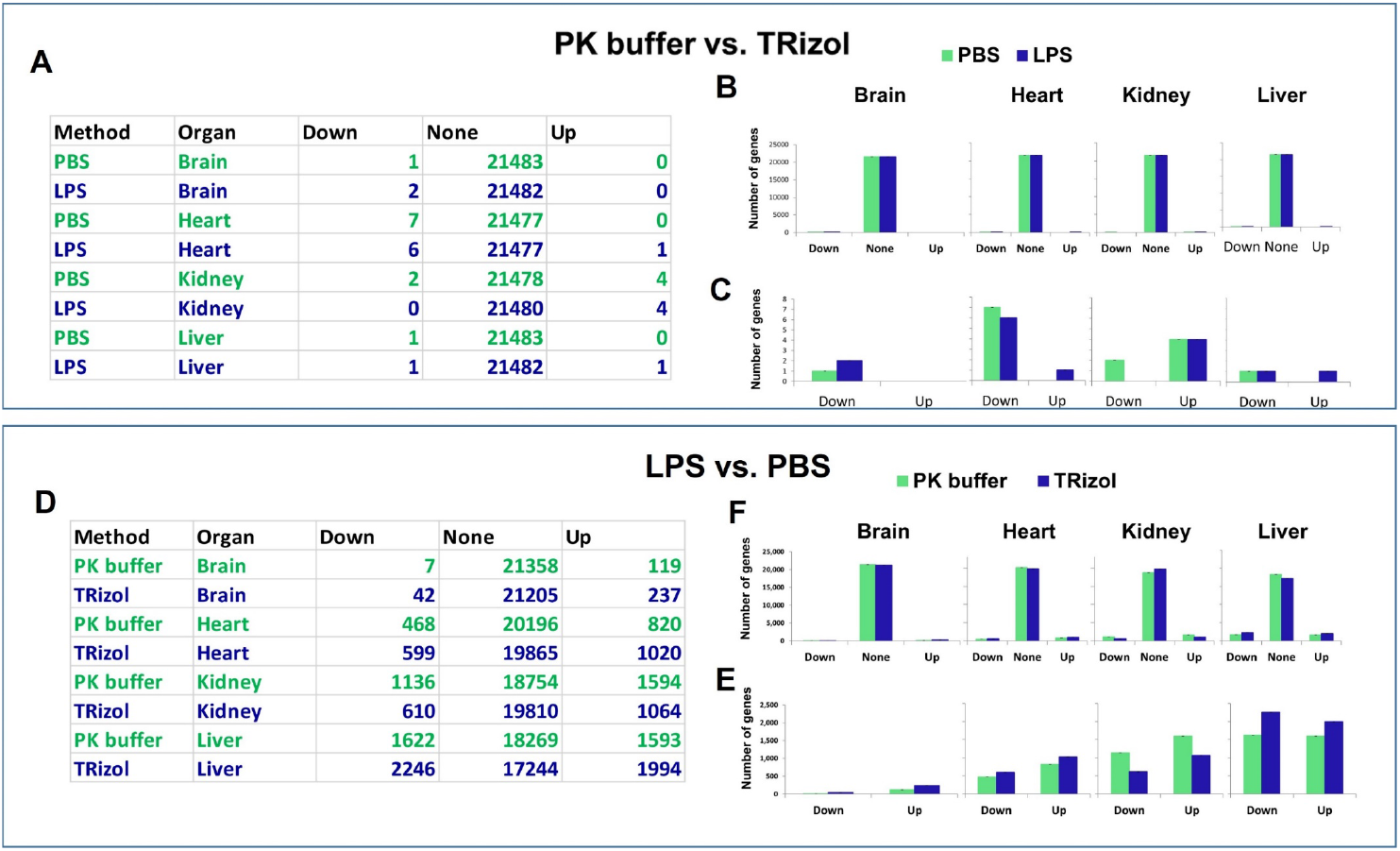
Numbers of differentially expressed genes in LPS endotoxin and PBS (control) treated mice. ***A-C*, PK buffer vs. TRIzol comparison.** *A*, table that shows the numbers for each group used to generate B-C plots. *B*, Comparison of the number of organ transcripts extracted with PK buffer (PBS) vs. TRIzol from either PBS (*green*) or LPS (blue) (y-axis) treated mice that were decreased (*down*), did not change (*none*) or increased (*up). C*, plot that shows the numbers of genes whose expression was downregulated or upregulated. ***D-E*, LPS vs. PBS comparison**. *D*, table that shows the numbers for each group used to generate E-F plots. *F*, Comparison of the number of organ transcripts extracted with either PK buffer (*green*) or TRIzol (blue) (y-axis) that were decreased (*down*), did not change (*none*) or increased (*up*) in response to LPS compared to PBS treatment. *E*, plot that shows the numbers of genes whose expression was downregulated or upregulated.

**Table S1.** Hardware and labware.

| Item Description | Manufacturer/Supplier | Location | Part No. |
| --- | --- | --- | --- |
| PIXUL™ 96-well sonicator | Matchstick Technologies | Kirkland, WA | P01199-01A |
|  | Active Motif | Carlsbad, CA | 53130 |
| CryoTray | Matchstick Technologies | Kirkland, WA | P01200-01A |
| CryoBlock | Matchstick Technologies | Kirkland, WA | P01201-01A |
| CryoCore | Matchstick Technologies | Kirkland, WA | P01202-01A |
| CryoCore trephines | Matchstick Technologies | Kirkland, WA | P01203-01A |
| PlateHandle | Matchstick Technologies | Kirkland, WA | P01204-01A |
| Fluke 52-II Dual Probe Thermometer | Fluke Corporation | Everett, WA | 674689 |
| Fluke 80PJ-1 Bead Probe | Fluke Corporation | Everett, WA | 750422 |
| Small Dual Angle Beta Radiation Shield (used as iPad stand) | Universal Medical Inc. | Oldsmar, FL | UM3400 |
| iPAD Pro | Apple | Cupertino, CA | Model A2377 |
| NextSeq2000 | Illumina | San Diego, CA | 20038897 |
| 4200 TapeStation | Agilent | Santa Clara, CA | G29991BA |
| Invitrogen™ Qubit™ 3 Fluorometer | ThermoFisher | Waltham, MA | Q33216 |
| NanoDrop 1000 Spectrophotometer | ThermoFisher | Waltham, MA |  |
| Eppendorf ThermoMixer® C | Eppendorf | Hamburg, Germany | 5382000023 |
| Costar 96-well polypropylene round-bottom (used in PIXUL) | Corning | Oneonta, NY | 3365 |
| GeneMate 96-well 0.2ml plate semi skirt | VWR | Radnor, PA | 490003-794 |
| MicroAmp Optical Adhesive Film (used in PIXUL) | Applied Biosystems | Carlsbad, CA | 4311971 |
| E1-ClipTip™ Electronic Multichannel Pipettes | ThermoFisher | Waltham, MA | 4671090 |
| 7900HT Fast Real-Time PCR System with 384-Well Block Module | ThermoFisher | Waltham, MA | 4329001 |

**Table S2.** Kits and enzymes.

| Item Description | Manufacturer/Supplier | Location | Part No. |
| --- | --- | --- | --- |
| Zymo-Seq RiboFree Total RNA Library Kit | Zymo Research | Tustin, CA | R3003 |
| Zymo RNA Clean & Concentrator-5 | Zymo Research | Tustin, CA | R1016 |
| SpeedBeads™ magnetic carboxylate modified particles | MilliporeSigma | Burlington, MA | GE65152105050250 |
| SuperScript IV Reverse Transcriptase | ThermoFisher | Waltham, MA | 18090050 |
| RNaseOUT™ Recombinant Ribonuclease Inhibitor | ThermoFisher | Waltham, MA | 10777-019 |
| RNase-Free DNase I | Lucigen | Middleton, WI | D9902K |
| Invitrogen™ Proteinase K, recombinant-Invitrogen | ThermoFisher | Waltham, MA | 25530015 |
| Standard Sensitivity Large Fragment Analysis Kit | Agilent | Santa Clara, CA | DNF-492 |
| High Sensitivity RNA ScreenTape Ladder | Agilent | Santa Clara, CA | 5067-5581 |
| High Sensitivity RNA ScreenTape Sample Buffer | Agilent | Santa Clara, CA | 5067-5580 |
| High Sensitivity RNA ScreenTape | Agilent | Santa Clara, CA | 5067-5579 |
| NextSeq™1000/2000 P2 | Illumina | San Diego, CA | 20046811 |
| Colibri™ Library Quantification Kit | Invitrogen | Carlsbad, CA | A38524500 |
| Quant-it dsDNA High-Sensitivity Assay Kit (Qubit) | Invitrogen | Carlsbad, CA | Q33120 |

**Table S3.**
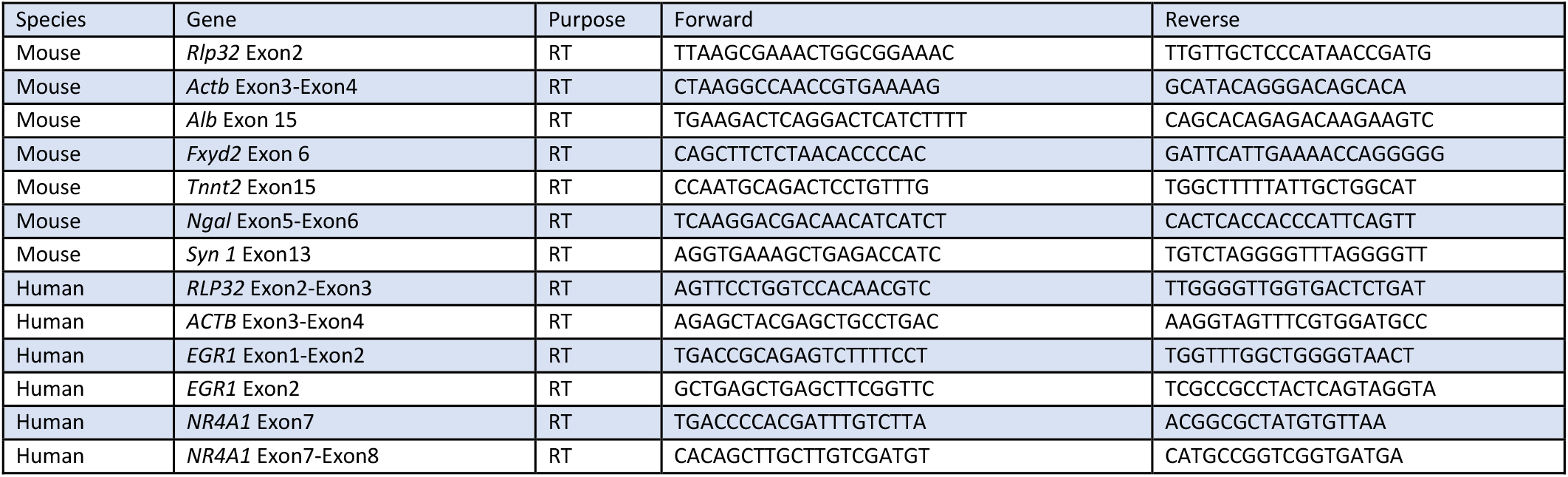
qPCR primers.

## References

1. Bomsztyk, K., Mar, D., Wang, Y., Denisenko, O., Ware, C., Frazar, C.D., Blattler, A., Maxwell, A.D., MacConaghy, B.E. and Matula, T.J. (2019) PIXUL-ChIP: integrated high-throughput sample preparation and analytical platform for epigenetic studies. Nucleic Acids Res, 47, e69.

2. Levy, S., Somasundaram, L., Raj, I.X., Ic-Mex, D., Phal, A., Schmidt, S., Ng, W.I., Mar, D., Decarreau, J., Moss, N. et al. (2022) dCas9 fusion to computer-designed PRC2 inhibitor reveals functional TATA box in distal promoter region. Cell Rep, 38, 110457.

3. Kanter, J.E., Shao, B., Kramer, F., Barnhart, S., Shimizu-Albergine, M., Vaisar, T., Graham, M.J., Crooke, R.M., Manuel, C.R., Haeusler, R.A. et al. (2019) Increased apolipoprotein C3 drives cardiovascular risk in type 1 diabetes. J Clin Invest, 129, 4165–4179.

4. Lim, M.D., Dickherber, A. and Compton, C.C. (2011) Before you analyze a human specimen, think quality, variability, and bias. Anal Chem, 83, 8–13.

5. Agrawal, L., Engel, K.B., Greytak, S.R. and Moore, H.M. (2018) Understanding preanalytical variables and their effects on clinical biomarkers of oncology and immunotherapy. Semin Cancer Biol, 52, 26–38.

6. Neumeister, V.M. and Juhl, H. (2018) Tumor Pre-Analytics in Molecular Pathology: Impact on Protein Expression and Analysis. Curr Pathobiol Rep, 6, 265–274.

7. Miescher, F. (1871) Ueber die chemische Zusammensetzung der Eiterzellen” (On the chemical composition of pus cells). Medicinisch-chemische Untersuchungen, 4, 441–460.

8. Thess, A., Hoerr, I., Panah, B.Y., Jung, G. and Dahm, R. (2021) Historic nucleic acids isolated by Friedrich Miescher contain RNA besides DNA. Biol Chem, 402, 1179–1185.

9. Hämmerling-J. (1953) Nucleo-cytoplasmic Relationships in the Development of Acetabularia. International Review of Cytology, 2, 475–498.

10. Temin, H.M. and Mizutani, S. (1970) RNA-dependent DNA polymerase in virions of Rous sarcoma virus. Nature, 226, 1211–1213.

11. Baltimore, D. (1970) RNA-dependent DNA polymerase in virions of RNA tumour viruses. Nature, 226, 1209–1211.

12. Mullis, K., Faloona, F., Scharf, S., Saiki, R., Horn, G. and Erlich, H. (1986) Specific enzymatic amplification of DNA in vitro: the polymerase chain reaction. Cold Spring Harb Symp Quant Biol, 51 Pt 1, 263–273.

13. Mullis, K.B. (1990) The unusual origin of the polymerase chain reaction. Sci Am, 262, 56-61, 64–55.

14. Chomczynski, P. and Sacchi, N. (2006) The single-step method of RNA isolation by acid guanidinium thiocyanate-phenol-chloroform extraction: twenty-something years on. Nat Protoc, 1, 581–585.

15. Chomczynski, P. and Sacchi, N. (1987) Single-step method of RNA isolation by acid guanidinium thiocyanate-phenol-chloroform extraction. Anal.Biochem., 162, 156–159.

16. Brown, R.A.M., Epis, M.R., Horsham, J.L., Kabir, T.D., Richardson, K.L. and Leedman, P.J. (2018) Total RNA extraction from tissues for microRNA and target gene expression analysis: not all kits are created equal. BMC Biotechnol, 18, 16.

17. Taryma-Lesniak, O., Sokolowska, K.E. and Wojdacz, T.K. (2020) Current status of development of methylation biomarkers for in vitro diagnostic IVD applications. Clin Epigenetics, 12, 100.

18. Ali, N., Rampazzo, R.C.P., Costa, A.D.T. and Krieger, M.A. (2017) Current Nucleic Acid Extraction Methods and Their Implications to Point-of-Care Diagnostics. Biomed Res Int, 2017, 9306564.

19. Slyper, M., Porter, C.B.M., Ashenberg, O., Waldman, J., Drokhlyansky, E., Wakiro, I., Smillie, C., Smith-Rosario, G., Wu, J., Dionne, D. et al. (2020) A single-cell and single-nucleus RNA-Seq toolbox for fresh and frozen human tumors. Nat Med, 26, 792–802.

20. Heid, C.A., Stevens, J., Livak, K.J. and Williams, P.M. (1996) Real time quantitative PCR. Genome Res, 6, 986–994.

21. Bomsztyk, K., Flanagin, S., Mar, D., Mikula, M., Johnson, A., Zager, R. and Denisenko, O. (2013) Synchronous Recruitment of Epigenetic Modifiers to Endotoxin Synergistically Activated Tnf-alpha Gene in Acute Kidney Injury. PLoS One, 8, e70322.

22. Andrews, S. (2010) FastQC: a quality control tool for high throughout sequence data.

23. Wang, L., Wang, S. and Li, W. (2012) RSeQC: quality control of RNA-seq experiments. Bioinformatics, 28, 2184–2185.

24. Ewels, P., Magnusson, M., Lundin, S. and Kaller, M. (2016) MultiQC: summarize analysis results for multiple tools and samples in a single report. Bioinformatics, 32, 3047–3048.

25. Liao, Y., Smyth, G.K. and Shi, W. (2019) The R package Rsubread is easier, faster, cheaper and better for alignment and quantification of RNA sequencing reads. Nucleic Acids Res, 47, e47.

26. Robinson, M.D., McCarthy, D.J. and Smyth, G.K. (2010) edgeR: a Bioconductor package for differential expression analysis of digital gene expression data. Bioinformatics, 26, 139–140.

27. Chen, Y., Lun, A.T. and Smyth, G.K. (2016) From reads to genes to pathways: differential expression analysis of RNA-Seq experiments using Rsubread and the edgeR quasi-likelihood pipeline. F1000Res, 5, 1438.

28. McCarthy, D.J., Chen, Y. and Smyth, G.K. (2012) Differential expression analysis of multifactor RNA-Seq experiments with respect to biological variation. Nucleic Acids Res, 40, 4288–4297.

29. Su, S., Law, C.W., Ah-Cann, C., Asselin-Labat, M.L., Blewitt, M.E. and Ritchie, M.E. (2017) Glimma: interactive graphics for gene expression analysis. Bioinformatics, 33, 2050–2052.

30. Monier B, McDermaid A, Zhao J and R, M.Q.v.C.r.v.o.R.d.i. (2021) vidger: Create rapid visualizations of RNAseq data in R. https://github.com/btmonier/vidger, https://bioconductor.org/packages/release/bioc/html/vidger.html.

31. Gillespie, M., Jassal, B., Stephan, R., Milacic, M., Rothfels, K., Senff-Ribeiro, A., Griss, J., Sevilla, C., Matthews, L., Gong, C. et al. (2022) The reactome pathway knowledgebase 2022. Nucleic Acids Res, 50, D687–D692.

32. Yu, G., Wang, L.G., Han, Y. and He, Q.Y. (2012) clusterProfiler: an R package for comparing biological themes among gene clusters. OMICS, 16, 284–287.

33. Gu, Z., Eils, R. and Schlesner, M. (2016) Complex heatmaps reveal patterns and correlations in multidimensional genomic data. Bioinformatics, 32, 2847–2849.

34. Harrel, F. (2021) Hmisc: Harrell Miscellaneous. https://CRAN.R-project.org/package=Hmisc.

35. Kim, D., Paggi, J.M., Park, C., Bennett, C. and Salzberg, S.L. (2019) Graph-based genome alignment and genotyping with HISAT2 and HISAT-genotype. Nat Biotechnol, 37, 907–915.

36. Danecek, P., Bonfield, J.K., Liddle, J., Marshall, J., Ohan, V., Pollard, M.O., Whitwham, A., Keane, T., McCarthy, S.A., Davies, R.M. et al. (2021) Twelve years of SAMtools and BCFtools. Gigascience, 10.

37. Ebeling, W., Hennrich, N., Klockow, M., Metz, H., Orth, H.D. and Lang, H. (1974) Proteinase K from Tritirachium album Limber. Eur J Biochem, 47, 91–97.

38. Wiegers, U. and Hilz, H. (1971) A new method using ‘proteinase K’ to prevent mRNA degradation during isolation from HeLa cells. Biochem Biophys Res Commun, 44, 513–519.

39. Gross-Bellard, M., Oudet, P. and Chambon, P. (1973) Isolation of high-molecular-weight DNA from mammalian cells. Eur J Biochem, 36, 32–38.

40. Lizardi, P.M. and Engelberg, A. (1979) Rapid isolation of RNA using proteinase K and sodium perchlorate. Anal Biochem, 98, 116–122.

41. Walsh, P.S., Metzger, D.A. and Higuchi, R. (1991) Chelex 100 as a medium for simple extraction of DNA for PCR-based typing from forensic material. Biotechniques, 10, 506–513.

42. Yu, J., Feng, Q., Ruan, Y., Komers, R., Kiviat, N. and Bomsztyk, K. (2011) Microplate-based platform for combined chromatin and DNA methylation immunoprecipitation assays. BMC Mol Biol, 12, 49.

43. Flanagin, S., Nelson, J.D., Castner, D.G., Denisenko, O. and Bomsztyk, K. (2008) Microplate-based chromatin immunoprecipitation method, Matrix ChIP: a platform to study signaling of complex genomic events. Nucleic Acids Res, 36, e17.

44. Blackburn, P. and Jailkhani, B.L. (1979) Ribonuclease inhibitor from human placenta: interaction with derivatives of ribonuclease A. J Biol Chem, 254, 12488–12493.

45. Ziegler, B.L., Lamping, C., Thoma, S. and Thomas, C.A. (1992) Single-cell cDNA-PCR: removal of contaminating genomic DNA from total RNA using immobilized DNase I. Biotechniques, 13, 726–729.

46. Kienzle, N., Young, D., Zehntner, S., Bushell, G. and Sculley, T.B. (1996) DNaseI treatment is a prerequisite for the amplification of cDNA from episomal-based genes. Biotechniques, 20, 612–616.

47. Winer, J., Jung, C.K., Shackel, I. and Williams, P.M. (1999) Development and validation of real-time quantitative reverse transcriptase-polymerase chain reaction for monitoring gene expression in cardiac myocytes in vitro. Anal Biochem, 270, 41–49.

48. Rumienczyk, I., Kulecka, M., Ostrowski, J., Mar, D., Bomsztyk, K., Standage, S.W. and Mikula, M. (2021) Multi-Organ Transcriptome Dynamics in a Mouse Model of Cecal Ligation and Puncture-Induced Polymicrobial Sepsis. J Inflamm Res, 14, 2377–2388.

49. Sharifian, R., Okamura, D.M., Denisenko, O., Zager, R.A., Johnson, A., Gharib, S.A. and Bomsztyk, K. (2018) Distinct patterns of transcriptional and epigenetic alterations characterize acute and chronic kidney injury. Sci Rep, 8, 17870.

50. Gharib, S.A., Mar, D., Bomsztyk, K., Denisenko, O., Dhanireddy, S., Liles, W.C. and Altemeier, W.A. (2016) System-Wide Mapping of Activated Circuitry in Experimental Systemic Inflammatory Response Syndrome. Shock, 45, 148–156.

51. Bomsztyk, K., Mar, D., An, D., Sharifian, R., Mikula, M., Gharib, S.A., Altemeier, W.A., Liles, W.C. and Denisenko, O. (2015) Experimental acute lung injury induces multi-organ epigenetic modifications in key angiogenic genes implicated in sepsis-associated endothelial dysfunction. Crit Care, 19, 225.

52. Remick, D.G., Ayala, A., Chaudry, I.H., Coopersmith, C.M., Deutschman, C., Hellman, J., Moldawer, L. and Osuchowski, M.F. (2019) Premise for Standardized Sepsis Models. Shock, 51, 4–9.

53. Earl, C.C., Smith, M.T., Lease, R.A. and Bundy, B.C. (2018) Polyvinylsulfonic acid: A Low-cost RNase inhibitor for enhanced RNA preservation and cell-free protein translation. Bioengineered, 9, 90–97.

54. Jacoli, G.G. (1968) Mechanism of inhibition of ribonuclease by bentonite. Can J Biochem, 46, 1237–1239.

55. Schroeder, A., Mueller, O., Stocker, S., Salowsky, R., Leiber, M., Gassmann, M., Lightfoot, S., Menzel, W., Granzow, M. and Ragg, T. (2006) The RIN: an RNA integrity number for assigning integrity values to RNA measurements. BMC Mol Biol, 7, 3.

56. Opitz, L., Salinas-Riester, G., Grade, M., Jung, K., Jo, P., Emons, G., Ghadimi, B.M., Beissbarth, T. and Gaedcke, J. (2010) Impact of RNA degradation on gene expression profiling. BMC Med Genomics, 3, 36.

57. Chen, E.A., Souaiaia, T., Herstein, J.S., Evgrafov, O.V., Spitsyna, V.N., Rebolini, D.F. and Knowles, J.A. (2014) Effect of RNA integrity on uniquely mapped reads in RNA-Seq. BMC Res Notes, 7, 753.

58. Kopczynski, M., Rumienczyk, I., Kulecka, M., Statkiewicz, M., Pysniak, K., Sandowska-Markiewicz, Z., Wojcik-Trechcinska, U., Goryca, K., Pyziak, K., Majewska, E. et al. (2021) Selective Extracellular Signal-Regulated Kinase 1/2 (ERK1/2) Inhibition by the SCH772984 Compound Attenuates In Vitro and In Vivo Inflammatory Responses and Prolongs Survival in Murine Sepsis Models. Int J Mol Sci, 22.

59. See, J.E. (2015) Visual Inspection Reliability for Precision Manufactured Parts. Hum Factors, 57, 1427–1442.

60. Okarma, K. (2020) Applications of Computer Vision in Automation and Robotics. Appl Sci-Basel, 10.

61. Shapiro, L.G. (2020) SURVEY Computer Vision: the Last 50 years. Int J Parallel Emergent Distrib Syst, 35, 112–117.

62. Byron, S.A., Van Keuren-Jensen, K.R., Engelthaler, D.M., Carpten, J.D. and Craig, D.W. (2016) Translating RNA sequencing into clinical diagnostics: opportunities and challenges. Nat Rev Genet, 17, 257–271.

63. Borisy, A.A., Elliott, P.J., Hurst, N.W., Lee, M.S., Lehar, J., Price, E.R., Serbedzija, G., Zimmermann, G.R., Foley, M.A., Stockwell, B.R. et al. (2003) Systematic discovery of multicomponent therapeutics. Proc Natl Acad Sci U S A, 100, 7977–7982.

64. Gemberling, M.P., Siklenka, K., Rodriguez, E., Tonn-Eisinger, K.R., Barrera, A., Liu, F., Kantor, A., Li, L., Cigliola, V., Hazlett, M.F. et al. (2021) Transgenic mice for in vivo epigenome editing with CRISPR-based systems. Nat Methods, 18, 965–974.

65. Singh, V.K. and Seed, T.M. (2021) How necessary are animal models for modern drug discovery? Expert Opin Drug Discov, 16, 1391–1397.

66. Cuklina, J., Pedrioli, P.G.A. and Aebersold, R. (2020) Review of Batch Effects Prevention, Diagnostics, and Correction Approaches. Methods Mol Biol, 2051, 373–387.

67. Bomsztyk, K. and Scheuerman, S. (2021) Tissue sample coring system. United States Patent Applicatiion Publication, US 2021/0404915A1.

68. Bomsztyk, K., Mar, D., Scheuerman, S. and Darlington, G. (2021) Tissue sample storing system. United States Patent Applicatiion Publication, US 2021/0386056A1.

